# A noncanonical GTPase signaling mechanism controls exit from mitosis in budding yeast

**DOI:** 10.1101/2024.05.16.594582

**Authors:** Xiaoxue Zhou, Shannon Y. Weng, Stephen P. Bell, Angelika Amon

## Abstract

In the budding yeast *Saccharomyces cerevisiae*, exit from mitosis is coupled to spindle position to ensure successful genome partitioning between mother and daughter cell. This coupling occurs through a GTPase signaling cascade known as the mitotic exit network (MEN). The MEN senses spindle position via a Ras-like GTPase Tem1 which localizes to the spindle pole bodies (SPBs, yeast equivalent of centrosomes) during anaphase and signals to its effector protein kinase Cdc15. How Tem1 couples the status of spindle position to MEN activation is not fully understood. Here, we show that Cdc15 has a relatively weak preference for Tem1^GTP^ and Tem1’s nucleotide state does not change upon MEN activation. Instead, we find that Tem1’s nucleotide cycle establishes a localization-based concentration difference in the cell where only Tem1^GTP^ is recruited to the SPB, and spindle position regulates the MEN by controlling Tem1 localization. SPB localization of Tem1 primarily functions to promote Tem1-Cdc15 interaction for MEN activation by increasing the effective concentration of Tem1. Consistent with this model, we demonstrate that artificially tethering Tem1 to the SPB or concentrating Tem1 in the cytoplasm with genetically encoded multimeric nanoparticles could bypass the requirement of Tem1^GTP^ and correct spindle position for MEN activation. This localization/concentration-based GTPase signaling mechanism for Tem1 differs from the canonical Ras-like GTPase signaling paradigm and is likely relevant to other localization-based signaling scenarios.

## INTRODUCTION

Small GTPases are key regulators in many cellular processes including cell signaling, actin organization, and membrane trafficking^1^. They typically function as molecular switches whose interaction with downstream effectors are regulated by changing nucleotide state^2^. Their high affinity for guanine nucleotides and low intrinsic GTP hydrolysis and GDP/GTP exchange activities underlie their success as molecular switches. Regulatory proteins such as GTPase activating proteins (GAPs) and guanine nucleotide exchange factors (GEFs) control these switches by modulating their GTP/GDP state in response to specific inputs^3^. A classic example is the founding member of the family, Ras, which regulates cell proliferation in response to growth factors^4^. Binding of growth factors to receptors at the cell surface leads to the recruitment of a GEF to the membrane, which promotes the conversion of membrane anchored Ras^GDP^ to Ras^GTP^. Ras^GTP^ then recruits and activates downstream effectors to signal proliferation. Although well established, it is unclear whether the same switch mechanism applies to all small GTPases involved in cell signaling.

The mitotic exit network (MEN) is a GTPase signaling cascade in the budding yeast *Saccharomyces cerevisiae* that regulates the exit from mitosis where the mitotic spindle disassembles, the nucleus divides, and chromosomes decondense. In budding cells, the site of cytokinesis or division plane is specified by the bud neck prior to mitosis^5^. To ensure proper genome partitioning, the mitotic spindle must be aligned along the mother-bud axis. The spindle position checkpoint (SPoC) blocks exit from mitosis until a spindle pole successfully enters the bud (Fig. 1A). This checkpoint is executed by establishing two zones in the cell with spatially distributed proteins (an inhibitory zone in the mother compartment and an activating zone in the bud)^6,7^ and a sensor, the MEN, which integrates multiple signals including spindle position to regulate cell cycle progression^8^ (Fig. 1A-B). Only when a spindle pole escapes the inhibitory zone in the mother and enters the activating zone in the bud in anaphase is the MEN activated, triggering mitotic exit. If the spindle is mis-positioned and remains entirely in the mother cell’s inhibitory zone, the MEN is kept inactive, arresting the cell in anaphase. Spindle position is transmitted to the MEN through the small GTPase Tem1^9,10^. How Tem1 senses and couples spindle position to MEN activation is not well understood.

**Figure 1:**
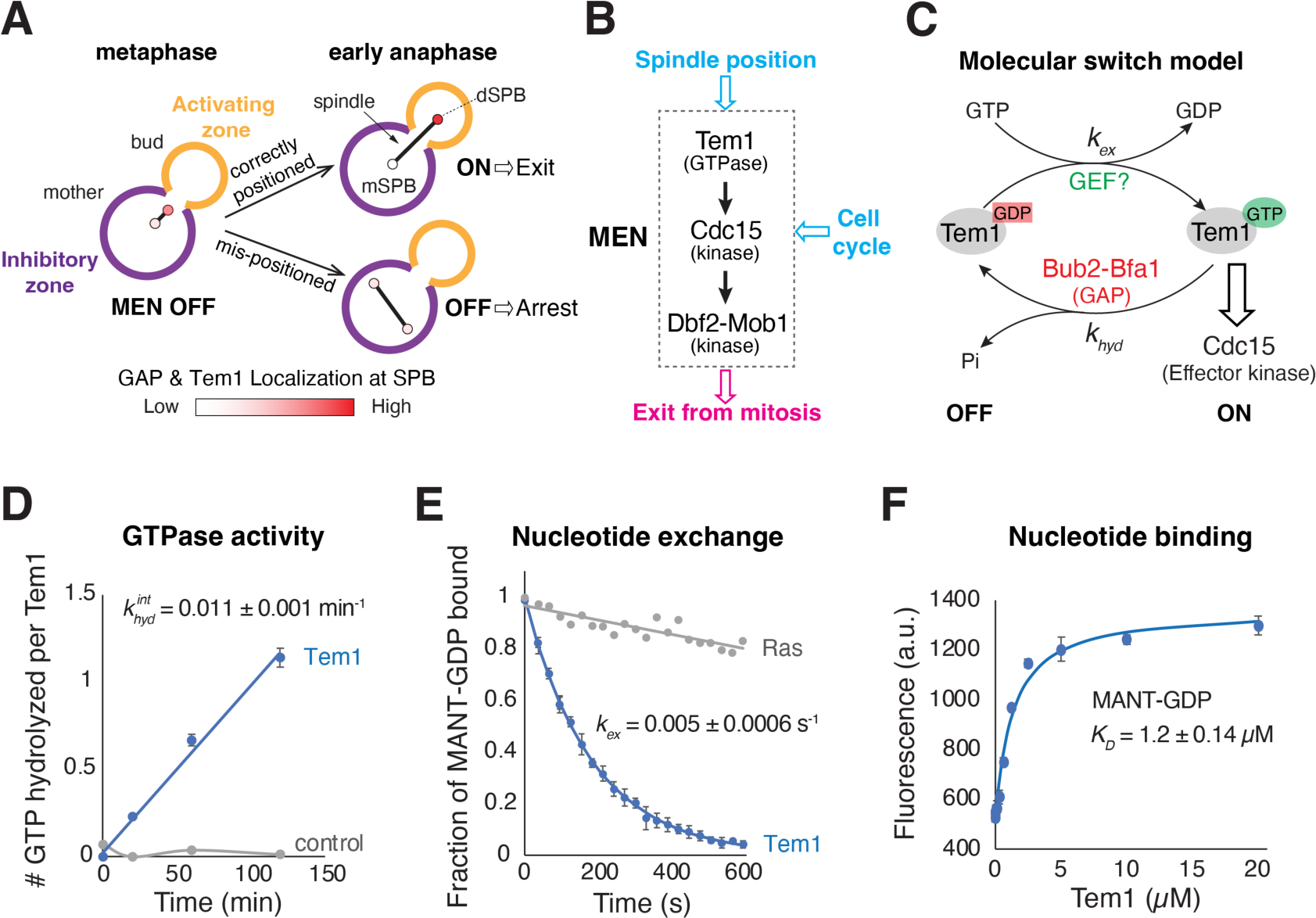
Tem1 is a noncanonical GTPase with relatively low affinity to nucleotides. **(A)** Illustration of the spindle position checkpoint in budding yeast and localization of the MEN GTPase Tem1 and its GAP. **(B)** Major components of the MEN and its inputs and outputs. **(C)** Molecular switch model for Tem1. **(D)** GTPase activity of Tem1. Recombinantly purified Tem1 was mixed with GTP and [α-^32^P]GTP on ice and the reaction was initiated by shifting to 30 °C. Samples were taken at indicated time points and analyzed by TLC (Fig. S1B-C). Shown are the average of 3 technical replicates with a linear fit. Error bars represent ± standard deviation (SD). **(E)** The rate of nucleotide exchange (MANT-GDP to GTP) for Tem1 and N-Ras. Recombinantly purified Tem1 (5 µM) and N-Ras (5 µM) were first incubated with MANT-GDP (200 nM) and the nucleotide exchange was initiated by the addition of excess GTP (1 mM). Shown are the average of 3 technical replicates with an exponential fit for Tem1 and a linear fit for N-Ras. Error bars represent ± SD. **(F)** The binding affinity of Tem1 for MANT-GDP. 200 nM MANT-GDP was titrated with increasing amount of purified apo-Tem1. Shown are the average from 3 technical replicates with a quadratic fit for the dissociation constant (*K_D_*). Error bars represent ± SD.

Like Ras, Tem1 undergoes a GTP/GDP nucleotide cycle and Tem1’s nucleotide states play an important role in its function. Mutations that block GTP hydrolysis and lock Tem1 in the GTP state are constitutively active and bypass the SPoC^11^. Conversely, nucleotide binding mutants that bias Tem1 toward the GDP state render the protein inactive and arrest cells in anaphase even with a correctly positioned spindle^12^. Given these findings, it has been assumed that Tem1 functions as a molecular switch like Ras and spindle position regulates Tem1 by changing its nucleotide state (Fig. 1C).

However, there are several aspects of Tem1 biology that are unusual for a small GTPase. First, while the GTP hydrolysis rate of Tem1 is accelerated by a bipartite GAP complex Bub2-Bfa1^10,13^, Tem1 does not appear to require a GEF to function, likely due to a relatively fast intrinsic nucleotide exchange rate^13^. As a result, the only way to regulate Tem1’s nucleotide state is by modulating the activity of the GAP. The polo-like kinase Cdc5 phosphorylates Bfa1^14^ which was shown to reduce Bub2-Bfa1 GAP activity in vitro^15^. Thus, a prevailing model is that the MEN is activated by inactivating the GAP^14–17^. A second peculiar aspect of Tem1 is that its cellular localization correlates with MEN activity^10,18,19^ (Fig. 1A). When the spindle is correctly positioned in anaphase (MEN ON), Tem1 primarily localizes to the spindle pole body (SPB, functional equivalent of centrosomes in yeast) that moves into the bud (daughter SPB or dSPB). However, if the spindle is mis-positioned (MEN OFF), Tem1 is largely cytoplasmic. Intriguingly, Tem1 localization to the SPB is highly dynamic with a residence time of around 5 s^17,18,20^. Furthermore, forcing Tem1 localization to the SPB by artificially tethering Tem1 to the SPB component Cnm67 overrides the checkpoint (SPoC defective)^20^, indicating a functional link between Tem1 localization and MEN activation. How Tem1’s nucleotide cycle and localization integrate to regulate MEN activation remains unclear.

Here, we demonstrate that Tem1 does not function as a molecular switch as its nucleotide state does not change upon MEN activation. Instead, Tem1’s GTP/GDP cycle is used to establish a localization-based concentration difference in the cell: only at the SPB can Tem1 achieve the effective concentration required to interact with its downstream effector protein Cdc15. Consequently, spindle position regulates MEN activation by controlling Tem1 localization. This noncanonical GTPase signaling mechanism differs from the molecular switch paradigm described by Ras and enables Tem1 to couple a spatial cue (i.e., spindle position) to downstream signaling. The observation that localizing a protein to SPB/centrosomes can modulate protein-protein interactions is likely relevant to other localization-based signaling scenarios.

## RESULTS

### Tem1 is a noncanonical small GTPase with relatively low affinity for nucleotides

To understand the role of Tem1’s nucleotide cycle, we first characterized the biochemical properties of purified Tem1 (Fig. S1): its intrinsic GTP hydrolysis and nucleotide exchange rates, and nucleotide binding activities. Tem1 hydrolyzed GTP with *k_hyd_* = 0.011 ± 0.001 min^−1^ (or 1.8 × 10^−4^ s^−1^, Fig. 1D) and exhibited a nucleotide exchange rate *k_ex_* = 0.005 ± 0.0006 s^−1^ (Fig. 1E). The hydrolysis rate is slower than those previously measured with an indirect method^13^, whereas the exchange rate is similar to prior reports^11,13^ and much faster than that of Ras (Fig. 1E). The fast intrinsic nucleotide exchange rate of Tem1 prompted us to examine whether Tem1 has a reduced affinity for nucleotides. Indeed, we found that Tem1 has a low micromolar affinity for a fluorescent derivative of GDP (MANT-GDP, *K_D_* = 1.2 ± 0.1 μM, Fig. 1F), in contrast to the picomolar to nanomolar nucleotide affinities for typical small GTPases^3^. These results suggest that while the intrinsic GTP hydrolysis rate is low for Tem1, the fast intrinsic nucleotide exchange renders Tem1 a self-activating molecular switch as the OFF state (Tem1^GDP^) is unstable due to the high GTP to GDP ratio in the cell^21^. Consequently, Tem1’s nucleotide state is mainly controlled by the GAP activity.

### Tem1’s effector protein Cdc15 has a weak preference for Tem1^GTP^

We next examined the interaction between Tem1 and its effector Cdc15. As an effector, we expected Cdc15 to have a strong preference for Tem1^GTP^. To measure the binding affinity between Cdc15 and Tem1 with different nucleotides, we developed an assay using AlphaLISA^22^, a bead-based technology that reports on biomolecular interactions. We mixed recombinant biotinylated Tem1 loaded with either GTPγS or GDP with Cdc15-eGFP from yeast lysate. Their interaction was detected with two populations of beads that bind Cdc15 and Tem1, respectively, and produce proximity-based luminescence upon interaction (Fig. 2A). Titrating biotinylated Tem1 promotes AlphaLISA signal until the amount of Tem1 exceeds the bead binding capacity, after which additional Tem1 competes with bead-bound Tem1 and reduces the signal (the hook point, Fig. 2B). This assay revealed that Cdc15 indeed had a higher affinity towards Tem1^GTPγS^ but also showed detectable binding towards Tem1^GDP^ prior to the hook point (Fig. 2B). Using a competition binding assay with unlabeled Tem1(Fig. 2D), we estimated the binding affinity of Cdc15 to Tem1^GTPγS^ (*K_D_* = 3.9 ± 0.8 nM) and Tem1^GDP^ (*K_D_* = 16 ± 3 nM).

**Figure 2:**
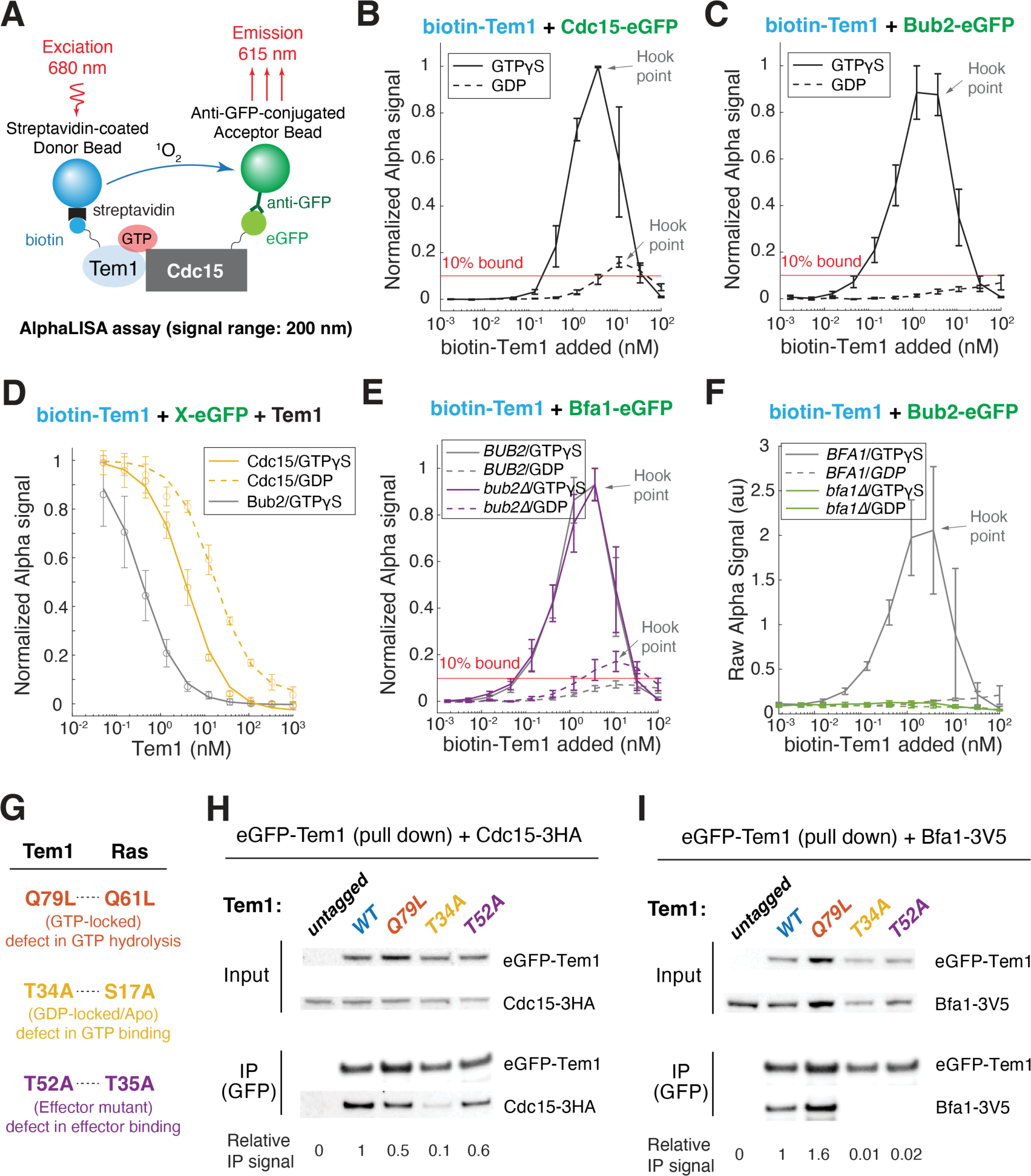
Tem1’s effector Cdc15 has a weak preference for Tem1^GTP^. (**A**) Illustration of the hybrid AlphaLISA assay used in this study. Recombinant biotinylated Tem1 were incubated together with cell lysate with eGFP tagged binding partners in the presence of GTPγS or GDP. The interaction was monitored with streptavidin coated Alpha donor beads and anti-GFP conjugated accepter beads. (**B-C**) Interaction between recombinant biotinylated Tem1 and eGFP tagged Cdc15 (B) or Bub2 (**C**) in lysate monitored by AlphaLISA. Diluted lysate of y1275 (Cdc15-eGFP) or y1378 (Bub2-eGFP) were incubated with different concentrations of recombinant biotinylated Tem1 in the presence of GTPγS or GDP. The Alpha signals were normalized to the maximum signal of each sample in the GTPγS condition. Black lines represent the average of three biological replicates and error bars denote standard deviation (SD). The Hook point is when the added biotin-Tem1 exceeds the binding capacity of the Alpha Donor beads which results in a competition between biotin-Tem1 in solution and on beads for binding. Red line indicates 10% of the available eGFP tagged protein (maximum binding capacity defined by the GTPγS condition) were bound by biotin-Tem1 on beads. (**D**) Dissociation constant of Tem1 binding to Cdc15 and the GAP estimated by a competition binding assay. Diluted lysate of y1275 (Cdc15-eGFP) or y1378 (Bub2-eGFP) were incubated with a fixed amount of biotinylated Tem1 and different concentrations of unlabeled Tem1 in the presence of GTPγS or GDP. Signal for Bub2-eGFP with GDP was too low to be quantified reliably. The Alpha signals were double normalized to the signal without competing Tem1 (normalized signal = 1) and without biotin-Tem1 (normalized signal = 0). Lines represent the average of three biological replicates and error bars denote SD. (**E**-**F**) Interaction between biotinylated Tem1 and eGFP tagged Bfa1 (E) or Bub2 (F). Diluted lysate of y1374 (Bfa1-eGFP), y1378 (Bub2-eGFP), y3099 (Bfa1-eGFP in *bub2Δ*), or y3632 (Bub2-eGFP in *bfa1Δ*) were incubated with different concentrations of biotinylated Tem1 in the presence of GTPγS or GDP, and the interaction was monitored with AlphaLISA. Lines represent the average of three biological replicates and error bars denote SD. **(G)** Point mutations generated in this study to modulate Tem1’s nucleotide state. (**H-I**) Interaction between different Tem1 mutants and the effector Cdc15 (H) or GAP complex (I) via co-immunoprecipitation. yEGFP-Tem1 was immunoprecipitated from exponentially grown cells of the indicated genotypes (y3461/y3452/y3472/y3457/y3456 or y2761/y2885/y2886/y3209/y3208) and the presence of Bfa1-3V5 or Cdc15-3HA were analyzed by Western blot analysis.

To validate our AlphaLISA results, we performed co-immunoprecipitation assays between Tem1 and Cdc15 in yeast lysates. To examine the effect of nucleotide state, we generated three mutations in Tem1 (Fig. 2G): a hydrolysis mutant, Tem1Q79L (corresponding to RasQ61L), that locks Tem1 in GTP state; a nucleotide-binding mutants, Tem1T34A (corresponding to Ras-S17A in the P loop region), that bias towards the GDP or apo state; and an effector mutant, Tem1T52A (corresponding to Ras-T35A at switch I region), that cannot undergo the confirmation change upon GTP binding necessary for effector recognition and thus essentially appears as “GDP-locked” for effectors. Contrary to classic GTPase effectors which show enhanced interaction with GTP-locked mutants and minimal interaction with GDP-locked and effector mutants, the GTP-locked Tem1Q79L precipitated less Cdc15 than wildtype Tem1 and the GDP-locked/apo Tem1T34A and effector mutant Tem1T52A retained considerable affinity for Cdc15 (≥10%, Fig. 2H, Fig. S2B). The observed weak nucleotide preference of Cdc15 is further supported by the AlphaFold2 predicted Cdc15-Tem1 interaction^23^ (Fig. S2C) where the binding interface is away from the nucleotide binding pocket of Tem1. We conclude that Cdc15 is a noncanonical GTPase effector protein with a relatively weak preference for Tem1^GTP^ over Tem1^GDP^. This unusual feature poses constrains on how well Tem1 could function as a molecular switch.

### The GAP complex Bub2-Bfa1 preferentially binds Tem1^GTP^

In contrast to the effector Cdc15, the GAP complex showed a strong preference for Tem1^GTPγS^ over Tem1^GDP^ with several orders of magnitude difference in affinity in the AlphaLISA assay (Fig. 2C, see 10% bound line): binding of the GAP to Tem1^GTPγS^ was readily detected even with < 0.1 nM biotinylated Tem1, whereas binding of the GAP to Tem1^GDP^ was barely detectable even with 100 nM Tem1. We estimated the binding affinity of the GAP complex to Tem1^GTPγS^ with the competition assay (Fig. 2D, *K_D_* < 0.4 ± 0.2 nM, ∼10-fold lower than Cdc15). We further dissected the contribution of each GAP subunit to the Tem1 interaction (Fig. 2E-F). Although Bub2 did not interact with Tem1 in the absence of Bfa1, Bfa1 bound similarly to Tem1^GTPγS^ in the presence or absence of Bub2. Interestingly, Bfa1 showed increased affinity for Tem1^GDP^ in the absence of Bub2 prior to the hook point (Fig. 2E), suggesting that Bfa1 alone and the Bub2-Bfa1 complex interact with Tem1 differently.

In parallel, we performed co-immunoprecipitation between different Tem1 nucleotide mutants and the GAP (Fig. 2I). As shown previously^11^ and consistent with our AlphaLISA results, the GTP-locked Tem1Q79L interacted more strongly with the GAP than wild-type Tem1 while the GDP-locked/apo mutant Tem1T34A and the effector mutant Tem1T52A had significantly reduced affinity for the GAP: almost no Bfa1 was pulled down by these Tem1 mutations (Figs. 2I & S2B). We conclude that the GAP complex specifically interacts with Tem1^GTP^. Based on their distinct nucleotide preference and sensitivity to mutations in Tem1 (Fig. S2B), our results also suggest that Cdc15 and the GAP complex do not share the exact same binding interface on Tem1.

### Tem1’s nucleotide state regulates its localization

How nucleotide state regulates Tem1 remains elusive. Given the correlation between Tem1’s SPB localization and its activity, we investigated whether Tem1 nucleotide state regulates its cellular localization. To track Tem1 localization in the cell, we developed a N-terminal fluorescent Tem1 fusion (yEGFP-Tem1) that, unlike previous C-terminal fusions^19,24^, showed similar expression and activity as wild-type untagged Tem1 (Fig. S3). Tem1 localizes primarily to the dSPB in metaphase and this localization is enhanced upon anaphase onset (which typically coincides with migration of the dSPB into the bud) followed by a decline as cells exit from mitosis (Fig. 3B). As previously reported^11^, the GTP-locked Tem1Q79L showed stronger localization to both SPBs throughout metaphase and anaphase (Fig. 3). In contrast, the inactive GDP-locked/apo mutants T34A abolished SPB localization and the effector mutant T52A significantly reduced SPB localization. These results suggest that Tem1’s nucleotide state specifies its SPB localization and that only Tem1^GTP^ is recruited to the SPB.

**Figure 3:**
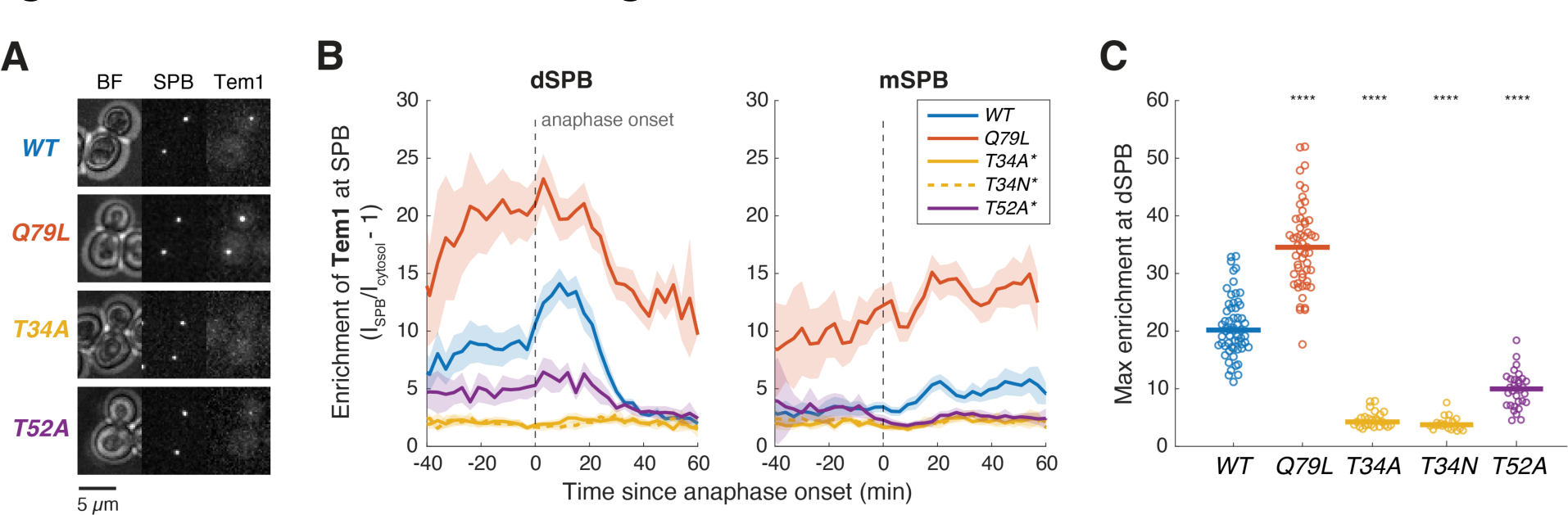
Tem1’s Nucleotide state regulates its localization. (**A-C**) Representative images and quantification of SPB localization in cells expressing yEGFP tagged wild-type Tem1 (y1748, *n* = 73 cells), GTP-locked hydrolysis mutant Q79L (y1824, *n* = 63 cells), GDP-locked/apo nucleotide binding mutant T34A/N (y3167/y3171, *n* = 29/21 cells), or effector binding mutant T52A (y3147, *n* = 32 cells) together with Spc42-mScarlet-I (SPB marker). Cells expressing *tem1T34A/N* and *tem1T52A* (marked with *) were kept alive with hyperactive *DBF2-HyA* that can rescue *tem1Δ*. Cells were grown at 25°C in SC medium + 2% glucose and imaged every 3 minutes for 4 hours. Single cell traces were aligned based on anaphase onset, as defined as spindle length > 3 μm (measured based on SPB marker Spc42-mScarlet-I), and averaged. Solid lines represent the mean, shaded areas represent 95% confidence intervals (B). For maximum enrichment at SPB (C), solid lines represent the median. \*\*\*\**p* < 0.0001 by two-sided Wilcoxon rank sum test for comparing with *WT*.

### SPB localization of Tem1 depends on the GAP Bub2-Bfa1 and effector Cdc15

How is Tem1^GTP^ recruited to the SPB? Previous studies showed that SPB localization of Tem1 mainly depends on the GAP complex Bub2-Bfa1^10^. Our finding that the GAP specifically interacts with Tem1^GTP^ could explain the nucleotide dependence of Tem1 localization. However, a small pool of Tem1 can localize to the SPB in a GAP-independent manner through an unknown mechanism. Using our yEGFP-Tem1 fusion and quantitative time-lapse microscopy, we compared the localization patterns of Tem1, the GAP component Bfa1, and Tem1’s effector protein Cdc15 in the presence or absence of the catalytic subunit of the GAP complex Bub2 (Fig. 4A).

**Figure 4:**
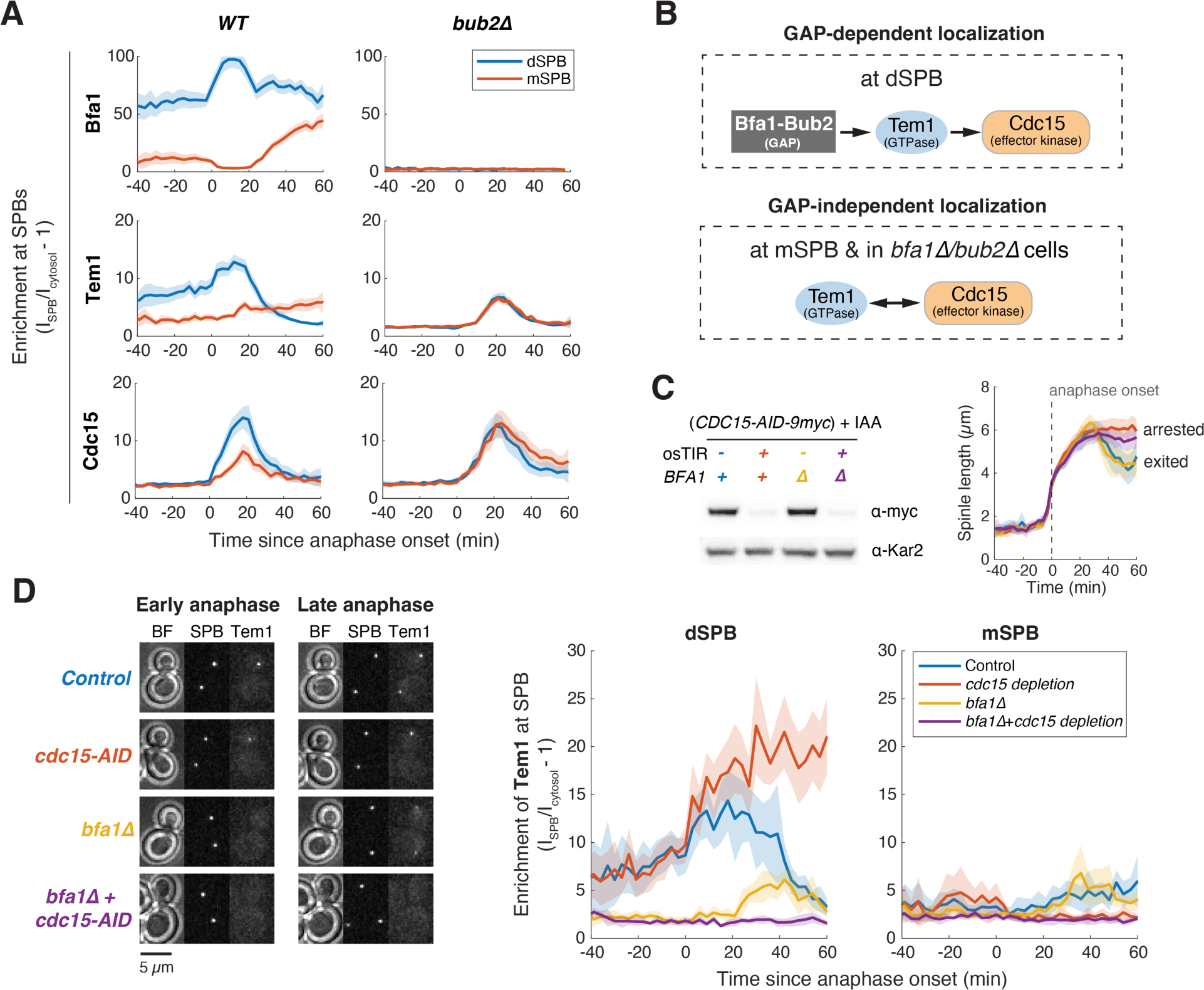
Localization of Tem1 to the SPB depends on its GAP and effector. **(A)** Comparison of SPB localization for Bfa1, Tem1 and Cdc15 with or without *BUB2*. Cells with the indicated *BUB2* alleles (columns) and yEGFP-labeled proteins (rows) were grown at 25 °C in SC medium + 2% glucose and imaged every 3 minutes for 4 hours. Strains imaged were y1374 (Bfa1-yEGFP, *n* = 104 cells), y3099 (Bfa1-yEGFP in *bub211*, *n* = 17 cells), y1748 (yEGFP-Tem1, *n* = 65 cells), y1929 (yEGFP-Tem1 in *bub211*, *n* = 72 cells), y1275 (Cdc15-yEGFP, *n* = 79 cells), and y1693 (Cdc15-yEGFP in *bub211*, *n* = 76 cells). **(B)** Dependency of SPB localization for two different modes of localization patterns observed. **(C)** Cdc15 was efficiently depleted with the auxin-inducible degron (*CDC15-AID*). Cells with the indicated genotypes (y3229 (*osTIR-*/*BFA1*), y3224 (*osTIR+*/*BFA1*), y3228 (*osTIR-*/*bfa111*), y3223 (*osTIR+*/ *bfa111*)) were grown at 25 °C in SC medium + 2% glucose + 200 μM IAA for 2 hours before harvesting for Western blot analysis (left). Anaphase progression (right) was monitored in these cells by measuring spindle length (distance between two SPBs). Cells with Cdc15 depleted arrested in anaphase with elongated spindles whereas non-depleted cells disassembled their spindle (as indicated by the shortening of the distance between SPBs in late anaphase) to exit from mitosis. **(D)** Representative images (left) and quantification (right) of Tem1 localization at the SPB with or without depleting Cdc15 (*CDC15-AID* +/− *osTIR*) in the presence or absence of *BFA1*. Cells of y3229 (Tem1 control, *n* = 16 cells), y3224 (Tem1 with Cdc15 depletion, *n* = 16 cells), y3228 (Tem1 in *bfa111*, *n* = 12 cells), y3223 (Tem1 in *bfa111* with Cdc15 depletion, *n* = 10 cells) were grown at 25 °C in SC medium + 2% glucose + 200 μM auxin IAA and imaged every 3 minutes for 4 hours. For all graphs, single cell traces were aligned based on anaphase onset, as defined in Fig. 3, and averaged. Solid lines represent the average. Shaded areas represent 95% confidence intervals.

Bub2 and Bfa1 form a stable complex that localizes to the SPB^10^. The GAP complex preferentially accumulates at the SPB closest to the bud (typically the dSPB) in metaphase. Upon anaphase onset, this asymmetry is exacerbated due to both an increase in localization at the dSPB and active removal of the GAP from the mSPB^17^. The latter outcome is achieved, at least in part, through phosphorylation of Bfa1 by the inhibitory zone protein Kin4^16,17^, a kinase that is restricted to the mother cortex and localizes to the mSPB in anaphase^25,26^. Tem1 localization to the SPB resembled the GAP until late anaphase, when Tem1 also localized to the mSPB (Fig. 4A, 12-20 min). The effector Cdc15 did not localize to the SPBs until anaphase onset and showed only a moderate bias toward dSPB. Bfa1 does not localize to either SPBs in the absence of Bub2^10^ (Fig. 4A). In contrast, Tem1 and Cdc15 showed synchronized and symmetric localization to both SPBs in late anaphase in *bub2Δ* cells (Fig. 4A, Fig. S4A). Similar localization patterns for Tem1 were found in *bfa1Δ* or *bfa1Δbub2Δ* double mutants (Fig. S4B-D). Furthermore, the GTP-locked Tem1Q79L showed a similar localization profile and anaphase progression as wild-type Tem1 in the absence of the GAP (Fig. S4C-D), consistent with our prediction that as a self-activating GTPase, Tem1 is mainly in the GTP-bound state without the GAP.

Given the correlated localization between Tem1 and Cdc15 in *bub2Δ* cells, we hypothesized that in the absence of the GAP, Tem1 (mainly Tem1^GTP^) and Cdc15 form a complex that is recruited to the SPB. In this model, Tem1 and Cdc15 localization to the SPB should be interdependent (Fig. 4B). It is well established that Cdc15 localization depends on Tem1^19^ (Fig. S5D), leading us to examine the role of Cdc15 in Tem1 localization: we either depleted Cdc15 with the auxin-inducible degron system^27^ (Fig. 4C-D) or generated *cdc15Δ* strains kept alive with a hyperactive *DBF2* allele^28^ (Fig. S5A-B). In the degron strains, Cdc15 depletion was nearly complete and arrested cells in anaphase (Fig. 4C). Tem1 localization to dSPB was not affected by Cdc15 depletion or deletion in *BFA1* cells but was completely dependent on Cdc15 in *bfa1Δ* cells (Figs. 4D, S5A-B). Similar results were observed for Tem1Q79L (Fig. S5C), demonstrating that Tem1 (Tem1^GTP^) cannot localize to the SPB on its own and relies on either the GAP or Cdc15 for recruitment to the SPB.

### GAP stimulated GTP hydrolysis drives the fast turnover of Tem1 at the dSPB

Because the GAP specifically recruits Tem1^GTP^ to the dSPB, we reasoned that the rapid turnover of Tem1 at the SPB observed previously^17,18,20^ could be a result of GAP-stimulated GTP hydrolysis and consequent dissociation of Tem1^GDP^. To test this hypothesis, we sought to compare the turnover of wild-type Tem1 and the hydrolysis deficient Tem1Q79L mutant at the dSPB (Fig. 5A) with fluorescence recovery after photobleaching (FRAP) analysis. We depleted Cdc15 to avoid potential interference of Cdc15-mediated SPB localization of Tem1, although we did not observe significant difference for Tem1 dynamics at the dSPB in anaphase cells with or without Cdc15 depletion (Fig. S6A). Inhibiting GTP hydrolysis with Tem1Q79L increased the residence time of Tem1 at the dSPB in anaphase cells from *t_1/2_* = 6 (± 2) s to *t_1/2_* = 16 (± 7) s (Fig. 5B). A similar reduction in Tem1 turnover was observed in cells with a catalytically dead GAP mutant *bub2R85A* (Fig. S6B). Thus, GAP-stimulated GTP hydrolysis drives the rapid turnover of Tem1 at the dSPB. Importantly, this finding allows us to use Tem1 residence time at the dSPB to estimate *in vivo* GAP activity as the difference of turnover rate between Tem1 and Tem1Q79L.

**Figure 5:**
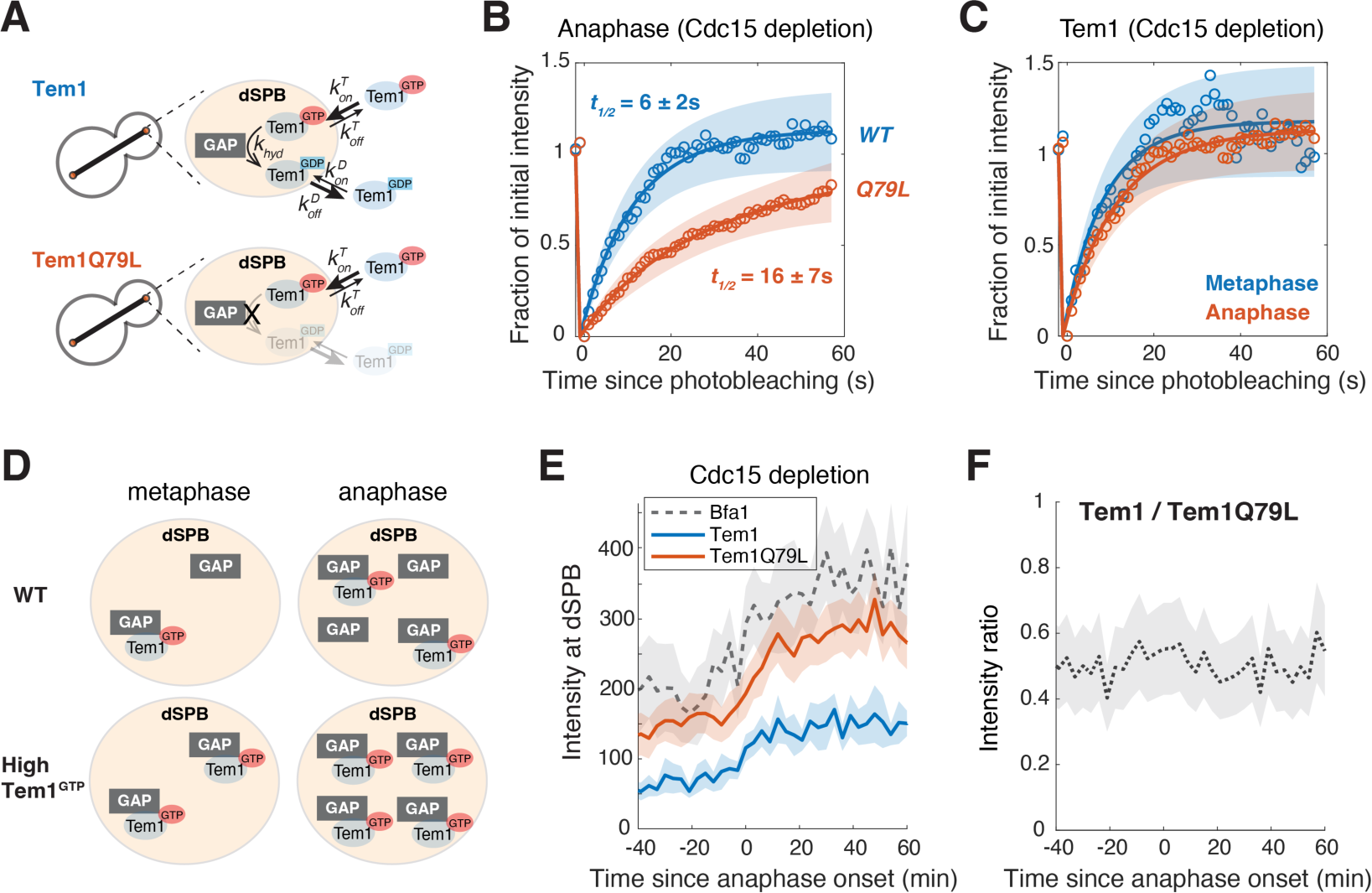
Tem1’s nucleotide state does not change upon MEN activation. **(A)** Illustration of the different routes for Tem1 and the GTP-locked Tem1Q79L at dSPB. Once Tem1^GTP^ is recruited to the dSPB by the GAP, its departure can occur either via Tem1^GTP^ dissociation (*k_off_^T^*) or GAP-stimulated GTP hydrolysis (*k_hyd_*) followed by dissociation (*k_off_^D^*). Assuming *k_off_^D^* >> *k_hyd_*, the residence time of Tem1 at the dSPB (*t_1/2_*^Tem1^) is set by the sum of two rates: *t_1/2_*^Tem1^ = ln2/(*k_off_^T^* + *k_hyd_*). For Tem1Q79L, its residence time (*t_1/2_*^Tem1Q79L^) is simply *t_1/2_*^Tem1Q79L^ = ln2/*k_off_^T^* assuming that Tem1^GTP^ and Tem1Q79L dissociate at similar rates. **(B)** FRAP analysis of Tem1 and TemQ79L at dSPB in anaphase. Cells of y3900 (*TEM1*, *n* = 9 cells) and y3903 (*TEM1Q79L*, *n* = 15 cells) were grown and imaged at room temperature in SC medium + 2% glucose + 100 μM IAA. Circles represent the average normalized fluorescence intensities after correcting for photo-bleaching during acquisition. Solid lines are the average fit and shaded areas represent SD. Half recovery time *t_1/2_* ± SD are indicated. **(C)** FRAP analysis of Tem1 at dSPB in metaphase and anaphase. Cells of y3900 were grown and imaged at room temperature in SC medium + 2% glucose + 100 μM IAA. **(D)** Proposed model for GAP and Tem1 localization at the SPB. Under wildtype condition (low [Tem1^GTP^] in the cell), only a fraction of the GAP complex at the dSPB is bound to Tem1^GTP^ in both metaphase and anaphase. While more GAP complex is recruited to dSPB in anaphase, the fraction of GAP bound to Tem1^GTP^ does not change between metaphase and anaphase. In cells with high [Tem1^GTP^] such as with *TEM1Q79L*, most of the GAP complex are bound to [Tem1^GTP^] because Tem1 is more abundant than the GAP. **(E)** dSPB localization of Tem1 (*WT* or *Q79L*) and Bfa1 in cells with Cdc15 depleted. Cells of y3900 (Tem1, *n* = 16 cells), y3903 (Tem1Q79L, *n* = 35 cells), and y4123 (Bfa1, *n* = 24 cells) were grown at 25 °C in SC medium + 2% glucose + 100 μM IAA and imaged every 3 minutes for 4 hours. Lines represent the average. Shaded areas represent 95% confidence intervals. **(F)** Ratio of dSPB intensities of Tem1 to Tem1Q79L from (E) was plotted to estimate the cellular level of Tem1^GTP^.

### The GAP is not inactivated in anaphase

It has been proposed that regulation of GAP activity, presumably by Cdc5 phosphorylation of Bfa1 at the SPB^15^, could regulate Tem1 in response to spindle position. Testing this hypothesis requires a way to assess Bub2-Bfa1’s GAP activity *in vivo*. We can now use Tem1 FRAP analysis to estimate the GAP-stimulated hydrolysis rate *k_hyd_* as the difference of SPB turnover rate between Tem1 and Tem1Q79L (*k_hyd_* = ln2/*t_1/2_*^Tem1^ - ln2/*t_1/2_*^Tem1Q79L^, Fig. 5A) which was 0.07 s^−1^. This is about 400-fold faster than Tem1’s intrinsic GTP hydrolysis rate, indicating that the GAP is active in anaphase.

If correct anaphase delivery of the dSPB to the bud downregulates GAP activity to promote Tem1 and MEN activation, then Tem1 residence time at the SPB should lengthen in anaphase. This, however, was not the case. Similar to previous reports^17,18,20^, we did not observe a significant difference in Tem1’s turnover rate at the SPB between metaphase and anaphase (Fig. 5C). As the turnover of Tem1 at dSPB is mainly driven by GAP-stimulated GTP hydrolysis (Fig. 5A-B), this result indicates that Bub2-Bfa1’s GAP activity at the SPB is not downregulated in anaphase compared to metaphase as previously suggested.

### Tem1’s nucleotide state does not change upon MEN activation

If the GAP is not inactivated in anaphase, does Tem1’s nucleotide state (ratio of Tem1^GTP^ to Tem1^GDP^ in the cell) change between metaphase and anaphase? Assuming that Tem1 undergoes spontaneous GEF-independent nucleotide exchange and GAP-stimulated GTPase hydrolysis, the parameters we measured experimentally combined with cellular concentrations of Tem1 and Bub2-Bfa1 from the literature^29^ (Fig. S7) allowed us to calculate that < 10% of Tem1 is in the GTP-bound state in the presence of active GAP, whereas 97% of Tem1 is in the GTP-bound state in the absence of the GAP (Fig. S7A). These ratios are predicted to stay the same between metaphase and anaphase given that GAP activity is not downregulated in anaphase.

To validate our conclusions, we sought to assess Tem1’s nucleotide state directly. Given that the GAP specifically recruits Tem1^GTP^ to the dSPB, we reasoned that the fraction of SPB localized GAP bound to Tem1 should correlate with cellular Tem1^GTP^ level ([Tem1^GTP^]) and thus can serve as an indicator for [Tem1^GTP^] (Fig. 5D). Indeed, the GTP-locked Tem1Q79L showed dramatically enhanced SPB localization (Fig. 3) due to high [Tem1^GTP^]. We quantified the dSPB localization of Bfa1, Tem1, and Tem1Q79L in cells with Cdc15 depleted (Fig. 5E). All three proteins increased dSPB localization upon anaphase onset synchronously due to the redistribution of GAP (Fig. S8). Since Tem1 is more abundant than the GAP complex (∼2.5x)^29^ and binds the GAP with sub-nanomolar affinity (Fig. 2D), we assumed that Tem1Q79L saturates all the GAP binding sites and could represent the maximum binding capacity. We thus estimated the fraction of GAP bound to Tem1 as the ratio of dSPB intensity of Tem1 to Tem1Q79L which remained steady over the cell cycle (Fig. 5F). These results strongly suggest that the nucleotide state of Tem1 (ratio of [Tem1^GTP^] to [Tem1^GDP^]) is not modulated to control MEN activation and the majority of Tem1 in the cell remains in the GDP-bound state in both metaphase (MEN inactive) and anaphase (MEN active).

### Localization of Tem1 to the SPB promotes MEN activation

If not by changing Tem1’s nucleotide state, how does spindle position regulate MEN activation? Given the functional link between SPB localization of Tem1 and MEN activation, we propose that spindle position regulates the MEN by controlling Tem1 localization. To test whether SPB localization of Tem1 directly controls MEN activation, we took advantage of the nucleotide binding mutants of Tem1 that fail to localize to SPB (Fig. 3). We hypothesized that these mutants are lethal due to mis-localization and artificially localizing them to the SPB should restore successful cell division. We tethered yEGFP-Tem1 to the SPB using a GFP nanobody (GFP binding protein, GBP) fused to a SPB outer plaque component Cnm67. This tethering completely rescued the lethality of both *tem1T34A/N* and *tem1T52A* (Fig. 6A). Direct fusion to Cnm67 also rescued an expanded panel of Tem1 mutants (Fig. S9). Furthermore, when Tem1 was tethered to the SPB, cells exited from mitosis with similar kinetics regardless of the nucleotide state (Fig. 6B). These results suggest that SPB localization of Tem1 is a key regulated step for MEN activation and that signaling downstream of Tem1 to the effector kinase Cdc15 does not require Tem1^GTP^ as long as Tem1 is correctly localized, consistent with the relatively weak nucleotide preference of Cdc15 (Fig. 2).

**Figure 6:**
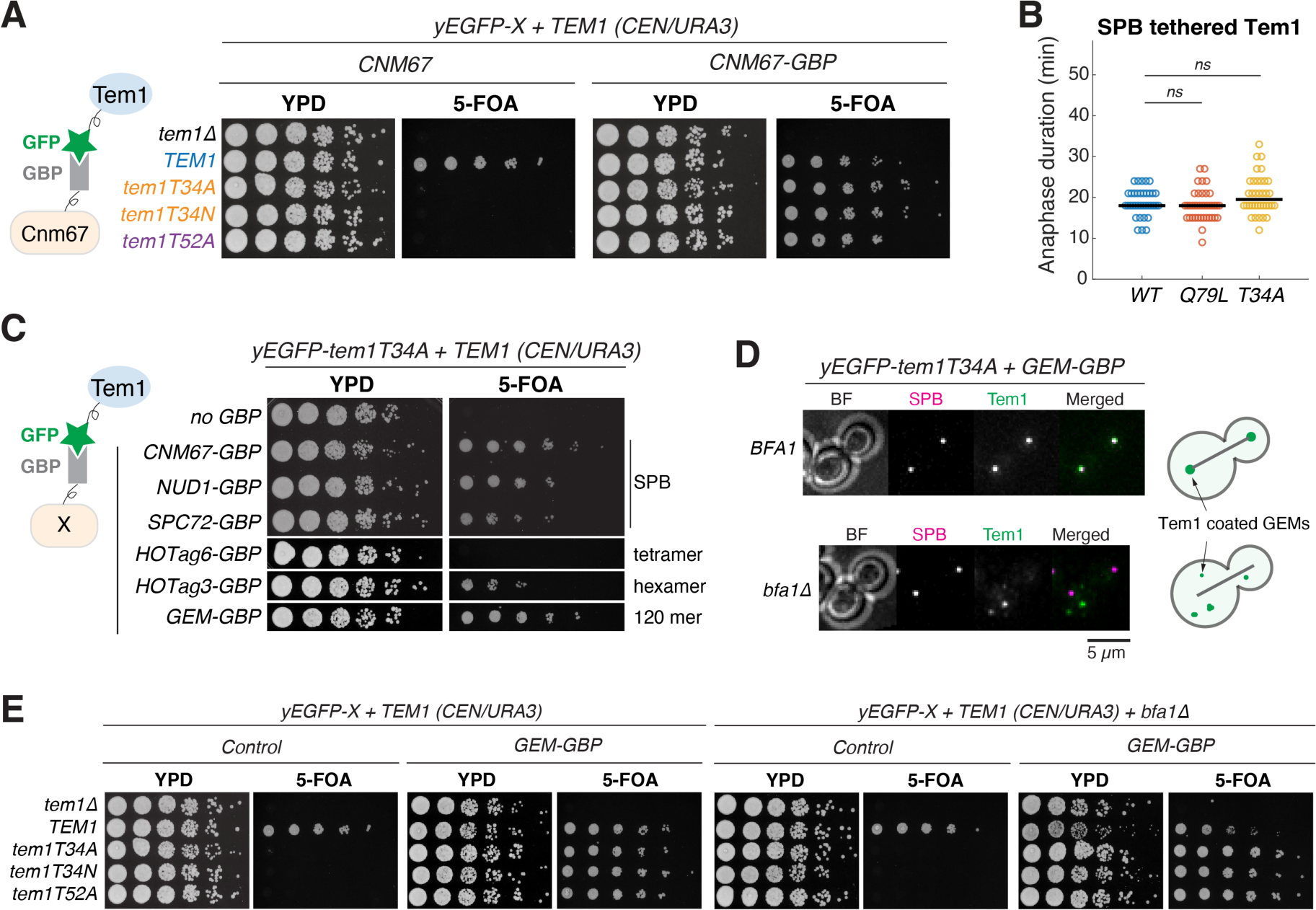
SPB localization modulates effective concentration of Tem1 for MEN activation. **(A)** Complementation analysis of Tem1 mutants with or without tethering to the SPB via *CNM67-GBP*. 5-fold serial dilutions of strains y2891/y3202/y3203/y3204/y3214 and y3303/y3304/y3305/y3306/y3310, with the indicated *TEM1* alleles, were spotted onto plates with or without 5’-fluoroorotic acid (5-FOA) and incubated at 25°C for 2 to 3 days. The presence of 5-FOA selects cells that are viable after losing the *TEM1 (URA3/CEN)* plasmid. GBP = GFP binding protein. **(B)** Distribution of anaphase duration for different *yEGFP*-*TEM1* alleles in the presence of *CNM67-GBP* (y3289, y3287, and y3291; *n* =46, 43, and 47 cells respectively). Cells were grown at 25°C in SC medium + 2% glucose and imaged every 3 minutes for 4 hours. Solid lines represent the median. **(C)** Complementation analysis of *tem1T34A* with and without tethering to different GBP fusion proteins. 5-fold serial dilutions of strains y3203/y3305/y3500/y3499/y3958/y3956/y3502 were spotted onto plates with or without 5-FOA and incubated at 25°C for 2 to 3 days. **(D)** Representative localization of GEM-tethered Tem1T34A in the presence or absence of *BFA1*. Cells of y3630 and y3557 were grown at 25 °C in SC medium + 2% glucose and imaged every 3 minutes for 4 hours. **(E)** Complementation analysis of different Tem1 mutants tethered to GEMs in the presence or absence of *BFA1*. 5-fold serial dilutions of strains y2891/y3202/y3203/y3204/y3214, y3970/y3973/y3502/y3976/y3985, y3971/y3974/y3948/y3977/y3986, and y3972/y3975/y3945/y3978/y3987 were spotted onto plates with or without 5-FOA and incubated at 25°C for 2 to 3 days.

### Concentrating Tem1 in the cytoplasm promotes MEN activation

As Cdc15’s key target Nud1^30^ is a core component of the SPB, the main function of Tem1 could be the recruitment of Cdc15 to the SPB. However, in *bubΔ* cells, MEN activation occurs even without the GAP-mediated recruitment of Tem1 to the SPB (Fig. 4A). We proposed that in the absence of the GAP, Tem1 and Cdc15 form a complex in the cytoplasm that can then localize to the SPB. To test whether the MEN can be activated by inducing Tem1-Cdc15 interaction in the cytoplasm, we modulated local concentrations of the localization-defective Tem1T34A by tethering it to homo-oligomers. Cdc15 forms dimers (or multimers)^31^ and due to an avidity effect could be sensitive to local concentrations of Tem1. Fusion to a tetrameric tether (HOTag6^32^) failed to rescue *tem1T34A* but fusion to a hexamer (HOTag3^32^) partially rescued the lethality (Fig. 6C). Tethering Tem1 to the surface of genetically-encoded multimeric nanoparticles (GEMs, 120mer)^33^ fully rescued the lethality of *tem1T34A* to the same degree as tethering to the SPB (Fig. 6C). These findings suggest that a high local concentration of Tem1 is sufficient for MEN activation.

Interestingly, Tem1T34A-coated GEMs clustered at the SPBs in a Bfa1-dependent manner (Figs. 6D & S10A), even though Tem1T34A did not localize to the SPB on its own (Fig. 3). This may be due to an avidity effect whereby the high density of Tem1T34A on GEMs compensated for the weak affinity of Tem1T34A for Bub2-Bfa1. To concentrate Tem1 away from the SPB, we repeated the GEM tethering experiment in *bfa1Δ* cells where the GEMs remained mostly in the cytoplasm (Fig. 6D & S10A). Under this condition, we observed the same rescuing effect for Tem1 lethal mutations defective in SPB localization (Fig. 6E). These results support a model in which SPB localization of Tem1 activates the MEN by modulating the effective concentration of Tem1 to promote Tem1-Cdc15 interaction and MEN signaling can be initiated in the cytoplasm by locally concentrating Tem1.

### SPoC is sensitive to Tem1 concentration

If spindle position regulates the MEN by modulating Tem1’s effective concentration, then increasing the global concentration of Tem1 should allow MEN activation even with a mispositioned spindle. To assess the sensitivity of SPoC to Tem1 concentration, we titrated cellular Tem1 level by introducing extra copies of the *TEM1* gene (Fig. S11A). We then quantified the integrity of the SPoC by depleting both Kar9 and Dyn1 using auxin-inducible degrons to trigger spindle mispositioning^7^ (Fig. 7A). Surprisingly, doubling Tem1 level with one extra copy (2x) was enough to compromise the SPoC (Fig. 7A, ∼40% SPoC defective). Cells with three extra copies (4x) were 95% SPoC defective (Fig. 7A). Furthermore, the time to exit mitosis in cells with mispositioned spindles decreased as Tem1 level was increased (Fig. 7B) revealing a dose-dependent relationship between Tem1 (Tem1^GTP^) level and MEN hyperactivation. Increasing cellular effective concentrations of Tem1 by local clustering instead of overexpression also bypassed the SPoC (Fig. S11C). Conversely, in cells expressing the GTP-locked Tem1Q79L mutant, lowering Tem1Q79L level with the auxin-inducible degron restored SPoC integrity (Fig. 7C, from 100% SPoC defective to ∼15%). These results demonstrate that SPoC is sensitive to cellular effective concentrations of Tem1 (Tem1^GTP^).

**Figure 7:**
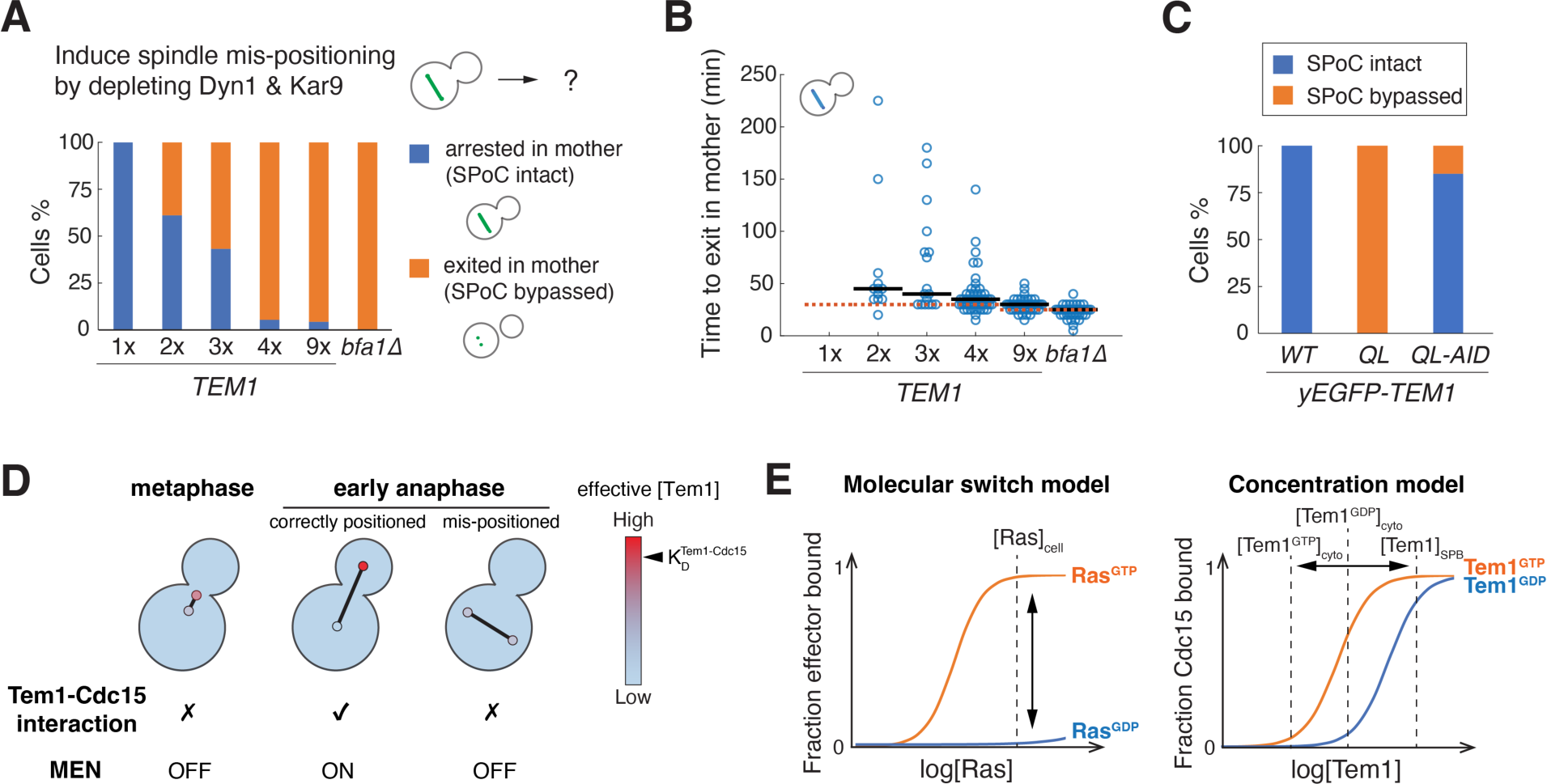
SPoC is sensitive to Tem1 concentration. (**A-B**) Evaluation of SPoC integrity in cells with different amount of Tem1. Cells of y3733 (1x*TEM1*, *n* = 102 cells), y3729 (2x*TEM1*, *n* = 72 cells), y3816 (3x*TEM1*, *n* = 64 cells), y3730 (4x*TEM1*, *n* = 102 cells), y3732 (9x*TEM1*, *n* = 101 cells), and y3815 (*bfa1τ1*, *n* = 65 cells) were grown at 25 °C in SC medium + 2% glucose + 100 μM IAA to induce spindle mispositioning by depleting Dyn1 and Kar9 and imaged every 5 minutes for 5 hours. Status of the spindle and cell cycle stages were monitored with GFP-Tub1. For time to exit in mother (mispositioned spindle disassembled in the mother cell) in D, solid lines represent the median, red dashed line indicate the median time to exit in bud (normal anaphase progression with correctly positioned spindle). (**C**) Evaluation of SPoC integrity in cells expressing Tem1Q79L. Cells of y3817 (yEGFP-*TEM1*, *n* = 55 cells), y3818 (yEGFP-*TEM1Q79L*, *n* = 47 cells), and y4311 (yEGFP-*TEM1Q79L-AID\**, *n* = 108 cells) were grown and monitored as described in (A). (**D**) Proposed model for Tem1 regulation. (**E**) Comparison between the canonical molecular switch model for Ras and the concentration model for Tem1.

## DISCUSSION

### Tem1 is a noncanonical GTPase that does not function as a molecular switch

We showed that Tem1 undergoes GEF-independent nucleotide exchange due to its lower binding affinity for guanine nucleotides (Fig. 1), and that Tem1’s effector protein Cdc15 has only a weak preference for Tem1^GTP^ (Fig. 2). This allows GDP-locked/apo Tem1 mutants that are localized to the SPB to activate the MEN. Most importantly, we demonstrated that Tem1’s nucleotide state ([Tem1^GTP^] : [Tem1^GDP^]) does not change upon MEN activation (Fig. 5). Our results suggest that Tem1 is regulated by a noncanonical mechanism where Tem1’s effective concentration is the key variable controlled by spindle position (Fig. 7D-E). Regulation of small GTPases by other noncanonical mechanisms like protein degradation (rather than changing nucleotide states) has been reported for the oncogene RIT1^34^ and the cytoskeleton regulator RhoB in quiescent endothelium^35^. To the best of our knowledge, Tem1 represents a novel example of small GTPases that functions via a non-switch mechanism regulated by changing localization.

### SPoC regulates the MEN by modulating effective Tem1 concentration

We propose a new model for Tem1 regulation and SPoC that relies on a localization-based concentration difference of Tem1 in the cell to regulate Tem1-Cdc15 interaction and MEN activation (Fig. 7D-E). In support of the model, perturbations that increase cellular effective concentrations of Tem1 either by overexpression (Fig. 7A), defective GTP hydrolysis (Fig. 7A-*bfa1Δ*, Fig. 7C-*TEMQ79L*), or local clustering (Fig. S11C) loosen the requirement of SPB localization of Tem1 for MEN activation and compromise the SPoC. Spindle position regulates localization of the Bub2-Bfa1 GAP to control Tem1 localization and concentration. This is achieved, at least in part, through the inhibitory zone protein Kin4 that actively removes the GAP complex from SPB(s) that remain in the mother compartment by phosphorylating Bfa1^16,17^. When the spindle is correctly positioned, the dSPB escapes the inhibitory mother compartment leading to the SPB localization of the GAP and Tem1 to trigger MEN activation. However, if the spindle is mispositioned and both SPBs remain in the mother cell, SPB localizations of the GAP and Tem1 are inhibited, keeping the MEN inactive. Decoupling SPB localization from spindle position by either tethering Tem1 or the GAP to the SPB also compromises the SPoC^11,17,20^. Finally, Tem1 is imported into the nucleus for degradation after exit from mitosis^36^ (Fig. S12) which facilitates MEN inactivation and resets Tem1 concentration for the following cell cycle.

It is still an open question exactly how Tem1-Cdc15 interaction leads to MEN activation. Cdc15’s kinase activity is not cell cycle regulated^37^. Instead, Cdc15 is thought to be mainly regulated by changing its localization and thus access to its target Nud1 at the SPB. Based on our finding of the GAP-independent SPB localization of Tem1 and Cdc15 (Fig. 4), we favor a model in which Tem1-Cdc15 complex formation promotes a form of active Cdc15 capable of localizing to the SPB. This could explain the previous finding that tethering Tem1 to the plasma membrane (PM) in the absence of the GAP inhibits MEN activation^20^. PM tethering sequesters Tem1 and thus Cdc15 and would prevent Cdc15’s access to Nud1. In contrast, when we tethered Tem1T34A to the GEMs, which are free to diffuse in the cytoplasm in the absence of the GAP, activated Cdc15 (presumably in complex with Tem1T34A on the surface of GEMs) localizes to the SPB (Fig. S10B-C) and brings Tem1T34A coated GEMs to the SPB as well (Fig. S10A).

### Additional cell cycle regulation on MEN activation

Our new model for Tem1 regulation explains how SPoC controls MEN activation in response to spindle position. However, Tem1 already localizes to the dSPB in metaphase when the MEN is supposed to be inactive, albeit at a lower level (∼50%) than in anaphase when the MEN is activated (Fig. 4A). Notably, Cdc15 does not localize to the SPB until anaphase (Fig. 4A). What is preventing Tem1 from interacting with Cdc15 and activating the MEN in metaphase? Given the sensitivity of MEN activation to Tem1 concentration (Fig. 7A), one possibility is that the two-fold concentration difference of SPB-localized Tem1 observed between metaphase and anaphase is enough to generate different outcomes for MEN activation. However, in diploid yeast with only one copy of *TEM1*, where SPB-localized Tem1 was reduced by ∼50% in anaphase (Fig. S11D), MEN activation was not prevented as these cells proceed to exit from mitosis with only a ∼3-min delay (Fig. S11D).

Another more plausible explanation is that there is an additional layer of regulation on Tem1-Cdc15 interaction that is cell-cycle dependent. It appears that in metaphase, SPB localized Tem1, although above the threshold concentration for Tem1-Cdc15 complex formation, is prevented from interacting with Cdc15. This inhibition is removed in anaphase to promote complex formation and MEN activation. As Tem1 is recruited to the SPB by Bub2-Bfa1 in metaphase (Fig. 4) and Bfa1 interacts with Tem1 on its own in a different manner than in the GAP complex (Fig. 2E), we propose that a likely candidate for the inhibitor of Tem1-Cdc15 interaction is Bfa1. In support of this model, overexpression of Bfa1 without its partner Bub2, arrests cells in anaphase^38–40^, presumably through inhibiting Tem1-Cdc15 interaction. Future experiments examining the specific binding interactions between Tem1 and Bfa1/Cdc15 will be required to test this hypothesis.

### The role of Tem1’s GTP/GDP cycle

Given that the key mode of regulation for Tem1 is by changing effective concentration, why does the system need a GTPase? In other words, what additional benefit might the GTP/GDP cycle provide for Tem1 regulation? Although Cdc15 has a relatively weak preference for GTP-bound Tem1, Tem1^GTP^ still interacts and activates Cdc15 much more efficiently. The GTP/GDP cycle thus provides an additional layer of concentration control to establish a concentration difference in the cell which prevents precocious MEN activation in the cytoplasm while remains primed for prompt MEN activation at the SPB. We found that in the absence of the GTP/GDP cycle with the GTP-locked Tem1Q79L, while it was possible to establish a semi-functional checkpoint by lowering the overall concentration of Tem1 (Fig. 7C), these cells took longer to exit from mitosis once the spindle is correctly positioned (Fig. S11E). Our results titrating Tem1 level (Figs. 7A-B, S11D) further indicates that with the GTP/GDP cycle *S. cerevisiae* cells operate with an optimum Tem1 concentration balancing the speed of mitotic exit and the necessary accuracy required for the checkpoint.

### Localization/concentration-based signaling mechanism

Our results support a model where SPB localization regulates Tem1 and MEN signaling by modulating Tem1’s effective concentration without changing its nucleotide state (Fig. 7D-E). The localization-based concentration difference of Tem1 could simply be attributed to high local concentration at the SPB and the avidity effect or an indirect mechanism where additional regulators at the SPB increase Tem1’s effective concentration. Our successful attempt in mimicking the concentration effect of SPB localization with the GEMs suggests that high local concentration is at least sufficient to initiate MEN signaling. The concentration effect we observed for Tem1 could be a general mechanism applicable to other localization-based signaling scenarios such as DNA damage response^41^ and kinetochore assembly^42^. We speculate that the same mechanism we described for Tem1 is conserved beyond budding yeast and could apply to other small GTPases. Future studies are needed to examine these possibilities.

## ACKNOWLEDGEMENTS

We thank E Unal (Berkeley, USA), J Haber (Waltham, USA), T Schwartz (Cambridge, USA), A Keating (Cambridge, USA), F Monje-Casas (Sevilla, Spain), S Piatti (Montpellier, France), and L Holt (New York, USA) for strains and reagents; A Murray and D Lew for their critical reading of the manuscript. This work was supported by National Institute of General Medical Science (K99GM140212 to XZ) and the Eunice Kennedy Shriver National Institute of Child Health and Human Development (HD085866 to AA). AA and SB were also investigators of the Howard Hughes Medical Institute.

## AUTHOR CONTRIBUTIONS

X.Z. and A.A. conceived the study. X.Z. performed all experiments. S.Y.W. generated Tem1 and Bub2 point mutants. X.Z. wrote the manuscript with input from S.P.B..

## MATERIALS AND METHODS

### Construction of Yeast strains and plasmids

All *Saccharomyces cerevisiae* yeast strains used in this study are derivatives of W303 (A2587) and are listed in Table S1. All plasmids used in this study are listed in Table S2. Yeast cells were cultured in standard YEP media (1% yeast extract, 2% peptone) with 2% D-glucose, or in standard Synthetic Complete media (SC) with 2% D-glucose. Cells were cultured at 25°C unless noted otherwise. C-terminal fusions and deletions were constructed using standard PCR-based methods. N-terminal tagging and point mutations of Tem1 and Bub2 were introduced at the endogenous locus using Cas9-mediated gene editing as described previously^43,44^. Successfully edited clones were checked by PCR and sequencing. Point mutations in GFP and small insertions in the tagging plasmids were introduced via Q5 site-directed mutagenesis. GFP and RFP (mCherry) nanobodies (GBP and RBP) were synthesized as gBlocks and inserted into tagging plasmids via Gibson assembly.

### Purification of Tem1 and biotinylated Tem1

Recombinant Tem1 was expressed as a 14xHis-SUMO fusion (p2579) in *Escherichia coli* Rosetta2(DE3) cells. Protein production was induced with 0.3 mM IPTG at 14 °C for 16-20 hours. Cells were collected by centrifugation at 6,000 x g, resuspended in lysis buffer (50 mM HEPES, pH 8.0, 500 mM NaCl, 2 mM MgCl_2_, 25 mM imidazole, 3 mM 2-mercaptoethanol (βME)) supplemented with cOmplete™ EDTA free protease inhibitor cocktail, treated with lysozyme for 30 mins at 4 °C followed by sonication. The lysate was cleared by centrifugation at 16,000 x g for 30 min. The soluble fraction was incubated with Ni-NTA agarose (QIAGEN) for 1 hour at 4 °C. The resin was washed with lysis buffer and transferred to low imidazole buffer (20 mM HEPES pH 8.0, 100 mM NaCl, 100 mM imidazole, 2 mM MgCl_2_) along with 2.5 μg SENP protease (gift from T. Schwartz’s lab^45^) and incubated at 4 °C overnight. Then the flow through containing the cleaved Tem1 was collected along with a 2 CV wash with low imidazole buffer and loaded onto a Mono S column equilibrated with buffer A (20 mM HEPES pH 8.0, 50 mM NaCl, 5 mM MgCl_2_, 10% glycerol, 1 mM DTT). After washing with buffer A, Tem1 was eluted with a linear gradient from 0 to 100% buffer B (20 mM HEPES pH 8.0, 1 M NaCl, 5 mM MgCl_2_, 10% glycerol, 1 mM DTT) in 20 CV. To make apo-Tem1, buffer EDTA (20 mM HEPES pH 8.0, 50 mM NaCl, 5 mM EDTA, 10% glycerol, 1 mM DTT) was used before elution to remove nucleotides bound to Tem1. Tem1 peak fractions were pooled and concentrated.

To generate biotinylated Tem1, Tem1 was expressed as a AviTag-6xHis fusion together with BirA (p3034, based on pDW363^46^) in *Escherichia coli* Rosetta2(DE3) cells. Protein production was induced with 0.5 mM IPTG at 16 °C in 1L LB supplemented with 15 mg biotin for 16-20 hours. Cells were collected and process as described above, and the soluble fraction was incubated with Ni-NTA agarose (QIAGEN) for 1 hour at 4 °C. The resin was washed with lysis buffer followed by low imidazole buffer (20 mM HEPES pH 8.0, 100 mM NaCl, 25 mM imidazole, 5 mM MgCl_2_) and biotinylated Tem1 was eluted with high imidazole buffer (20 mM HEPES pH 8.0, 100 mM NaCl, 250 mM imidazole, 5 mM MgCl_2_) and loaded onto a Mono S column for additional purification (see above). Tem1 peak fractions were pooled and concentrated.

### GTPase assay

GTP hydrolysis was measured for recombinant Tem1 with [α-^32^P]-GTP. Tem1 (3.3 µM final concentration) was mixed with GTP (33 µM final) supplemented with 5 µCi [α-^32^P]-GTP (PerkinElmer) in the reaction buffer (20 mM HEPES pH 7.5, 25 mM NaCl, 5 mM MgCl_2_) on ice and the reaction was initiated by shifting to 30 °C. At indicated time intervals, 2 µl reaction was removed and quenched with quench buffer (50 mM EDTA, 2% SDS) and analyzed via thin layer chromatography (TLC) on PEI-cellulose plates (Millipore sigma 105579). The plates were developed in 0.75M KH_2_PO_4_ (pH 3.5) and dried plates were exposed to storage phosphor screens overnight and scanned on a Typhoon phosphorimager (Cytiva). It is worth noting that when optimizing assay conditions, we found that EDTA which is often included in GTPase assays for small GTPase to increase nucleotide exchange or turnover did not make a difference in the rate of GTP hydrolysis for Tem1, consistent with Tem1’s high intrinsic nucleotide exchange rate.

### Nucleotide exchange and binding assay

To load Tem1 with fluorescently labeled nucleotides (i.e., MANT-GDP) for nucleotide exchange assays, recombinant Tem1 was first diluted in Tem1 dilution buffer (20 mM HEPES pH 7.5, 100 mM NaCl, 5mM MgCl_2_, 10% glycerol, 0.2 mg/ml BSA, 2 mM DTT) to 10 µM and mixed with equal volume of 400 nM MANT-GDP or MANT-GTP/GTPγS in Tem1 binding buffer (20 mM HEPES pH 7.5, 100 mM NaCl, 5mM MgCl_2_). The mixture was incubated for at least 10 minutes at room temperature to reach equilibrium and then kept on ice. Three technical replicates of 18 µl MANT-GDP/GTP/GTPγS loaded Tem1 mixture were plated in a black 384-well nonbinding microplate (Greiner Bio-One) and read in a SpectraMax ID5 plate reader (Molecular Dynamics) at 30 °C. To initiate nucleotide exchange, 2 µl of 10 mM unlabeled nucleotide (i.e., GTP) was added (1 mM final concentration) and readings were taken at 30s interval with 355 nm excitation and 448 nm emission. *k_obs_* was measured by fitting a single exponential decay after background subtraction.

To measure binding affinities, apo-Tem1 was diluted with Tem1 dilution buffer first to 40 µM then 2-fold serial diluted 10 times with the dilution buffer. Equal volumes of 400 nM MANT-GDP or MANT-GTPγS in Tem1 binding buffer were mixed with the apo-Tem1 dilution series and 15 µl of each mixture was placed in a black 384-well nonbinding microplate (Greiner Bio-One) and read in a SpectraMax ID5 plate reader (Molecular Dynamics) after 30 minutes incubation at 30 °C. Three independent dilutions were performed for each nucleotide conditions. *K_D_* was measured by fitting a quadratic equation: 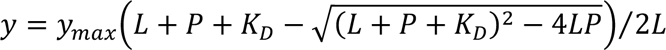, where *y_max_* is the maximum fluorescence signal at saturated binding, *L* is the total ligand concentration (MANT-GDP/GTPγS, 0.2 µM), *P* is total Tem1 concentration.

### Microscopy and image analysis

For live-cell microscopy, cells were imaged on agarose pads (2% agarose in SC medium + 2% glucose, unless otherwise noted) affixed to a glass slide and covered with a coverslip. Imaging was performed on a DeltaVision Elite platform (GE Healthcare Bio-Sciences) with an InsightSSI solid state light source, an UltimateFocus hardware autofocus system and a model IX-71, Olympus microscope controlled by SoftWoRx software. A 60x Plan APO 1.42NA objective and CoolSNAP HQ2 camera were used for image acquisition. For each time point, 8 *z* sections with 0.75 μm spacing were collect for each channel and were deconvolved. Maximum projections of the deconvolved *z* stack were used for fluorescence quantification.

Image analysis was performed with custom scripts in MATLAB as described previously. First, yeast cells were segmented and tracked through time using the bright-field image stacks. Next, fluorescence images of cell cycle markers (such as Spc42 or NLS) were segmented and tracked based on cell segmentation. Appearance of a cell cycle marker in a cell during the acquisition period was used to identify buds (daughter cells) and cell division events. Tracking of the cell cycle markers that migrated into the buds were used to identify the corresponding mother cells. Finally, for each division event identified, localization of the target protein at regions defined by the cell cycle markers was quantified.

For localization at the SPBs (*I_SPB_*), maximum intensity of the target protein at SPBs (based on segmentation of Spc42) was used given that the size of SPBs (∼100 nm) is within the diffraction limit of light microscopy. To calculate the relative enrichment of target protein at the SPB, intensities were normalized to the median intensity in the cytosol: *I_SPB_*⁄*I_cytosol_* − 1. Single cell traces were aligned based on the timing of anaphase onset or the movement of a SPB into the bud as indicated in figures, and averaged. 95% confidence intervals were calculated as *μ* ± 1.96 ∗ *σ*⁄√*n*, where *μ* and *σ* denote the mean and standard deviation respectively and *n* is the number of cells measured.

### Spotting assays

Tem1 mutants were constructed in the native locus with CRISPR in y3059 carrying a hyperactive Dbf2. Correct mutants were then crossed with y2892 (*tem1* knockout carrying a URA3-based CEN covering plasmid expressing wild-type Tem1) to generate strains with Tem1 mutants and the covering plasmid. Additional alleles tested were also introduced via cross. For spotting assays, these strains were grown in YPD overnight and diluted to OD 0.2 in YPD and grow for at least one doubling at room temperature. 5-fold serial dilutions were made for each culture starting from OD 0.2 and 4 µl were spotted onto YPD or plates with 5’-fluoroorotic acid (5-FOA) which selects cells that are viable after losing the URA3 covering plasmid.

### Immunoblot analysis

Log-phase cultures of cells grown in YEP + 2% glucose were harvested and treated with 5% TCA at 4°C overnight. TCA treated cell pellets were washed with acetone, air dried, and resuspended in lysis buffer (10 mM Tris, 1 mM EDTA, 2.75 mM DTT, pH = 8). Cells were lysed by bead-beating using a Multivortexer (max speed, 20 minutes) and glass beads at 4°C and followed by boiling in SDS PAGE protein loading buffer for 5 minutes. Lysates were clarified by centrifugation and were resolved with NuPAGE 4-12% Bis-Tris protein gel (Thermo Fisher Scientific) prior to transfer onto nitrocellulose membranes. Tem1 was detected with an anti-Tem1 antibody generated in rabbit against the peptide CKKLTIPEINEIGDPLLIYKHL^24^ at a 1:300 dilution. GFP was detected using an anti-GFP antibody (Clontech, JL-8) at a 1:1000 dilution. Myc tags were detected using an anti-Myc antibody (abcam, 9E10) at a 1:500 dilution. V5 tags were detected using an anti-V5 antibody (Invitrogen) at a 1:2000 dilution. HA tags were detected with an anti-HA.11 antibody (BioLegend) at 1:1000 dilution. Clb2 was detected using a rabbit anti-Clb2 at 1:1000 dilution. Kar2 was detected using a rabbit anti-Kar2 antiserum at a 1:200,000 dilution. DyLight 800 conjugated secondary antibodies (Cell Signaling) were used at a 1 :10,000 dilution. Blots were imaged using the ChemiDoc MP imager system (BioRad).

### Co-immunoprecipitation analysis

Log-phase cultures (∼ 40 OD of cells) were harvested and snap frozen in liquid nitrogen. Cell pellets were thawed in lysis buffer (50 mM Tris pH=7.5, 150mM NaCl, 1% NP40, 5mM MgCl2, 1mM DTT, supplemented with cOmplete™ EDTA free protease inhibitor and PhosSTOP™ inhibitor) and lysed via bead-beating with silica beads in a FastPrep-24™ (MP Biomedicals). Lysates were clarified by centrifugation and protein concentration in the lysates was measured with Bradford assay (Bio-Rad). 15 µl of GFP-Trap agarose (ChromoTek) was washed three times with NP40 buffer (50 mM Tris pH=7.5, 150mM NaCl, 1% NP40, 5mM MgCl2) and incubated with clarified lysates containing ∼3 mg of total protein per sample for 1 hour at 4 °C. The agarose beads were washed five times with NP40 buffer. Afterwards, the bound proteins were eluted by boiling the beads in protein loading buffer and resolved on NuPAGE 4–12% Bis-Tris protein gels (Thermo Fisher Scientific) followed by immunoblot analysis (see above).

### AlphaLISA analysis

Purified biotinylated Tem1 was first diluted to 500 nM and incubated with 100 µM GTPγS or GDP in assay buffer (50 mM Tris pH=7.5, 150mM NaCl, 1% NP40, 5mM MgCl2, 1mM DTT, 1x cOmplete™ EDTA free protease inhibitor and PhosSTOP™ inhibitor, 0.1% BSA) and 3-fold serial diluted. Equal volume of 100 µg/ml (5x final concentration) Alphascreen Streptavidin Donor beads (PerkinElmer 6760002S) diluted in assay buffer were added and incubated with nucleotide loaded Tem1 for 1 hour at room temperature in the dark. Log-phase cultures (∼ 10 OD of cells) containing eGFP tagged Bub2/Bfa1/Cdc15 were harvested and snap frozen in liquid nitrogen. Cell pellets were processed as described for co-immunoprecipitation assays and diluted to 3 mg/ml total protein concentration with lysis buffer. 5 µl lysate was added to each 384 well Alpha plates (PerkinElmer) together with 10 µl Tem1 coated donor beads and 10 µl 50 µg/µl AlphaLISA Anti-GFP Acceptor Beads (PerkinElmer AL133C) diluted in assay buffer. The plate was incubated for 4-6 hours at 4 °C and read using a M1000 plate reader (Tecan) with the default AlphaScreen settings at room temperature. Alpha signals were analyzed by first subtracting background signal measured with control yeast lysate without eGFP (y1584) and then normalized with the maximum signal of each experiment. For the competition assays with unlabeled Tem1, 0.5 nM and 5 nM final concentration of biotinylated Tem1 were used for the GTPγS and GDP binding assays respectively.

### FRAP analysis

FRAP analysis was performed on a DeltaVision-OMX Super-Resolution Microscope (Applied Precision, GE Healthcare Bio-Sciences) using a 60x oil objective and a 488 nm laser adjusted to bleach an area of approximately 0.5 μm in radius. Two prebleach images were acquired followed by a laser pulse (50% intensity) of 0.02 s duration and postbleach images were acquired at 1 s/frame for 60 s. Images at each time points were maximum projections of 5 *z* stacks with 0.5 μm spacing. Images were analyzed with a custom MATLAB script. After subtracting the background, fluorescence intensities in the cytosol, *I_cytosol_* (*t*), and at the SPB, *I_SPB_* (*t*) were measured after segmenting the cell and SPBs. Photobleaching was corrected by normalizing *I_SPB_* (*t*) with *I_cytosol_* (*t*), *I_SPB_norm_* (*t*) = *I_SPB_* (*t*) / *I_cytosol_* (*t*). Double normalization for FRAP was calculated to scale the photobleaching effect between 0 and 1: *I_SPB_FRAP_* (*t*) = [*I_SPB_norm_* (*t*) - *I_SPB_norm_* (*0*)] / [*I_SPB_norm_* (*pre*) - *I_SPB_norm_* (*0*)], where *t* = 0 is the time point (frame) right after photobleaching and *t* = *pre* is the time point (frame) right before photobleaching. The double normalized FRAP curves were then fitted to a single exponential curve: 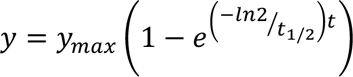 is the fraction recovered while *t*_4/%_ is the half-recovery time.

### SPoC assay

Spindle mispositioning was induced by depleting both Kar9 and Dyn1 with the auxin-inducible degron system. Spindle position and checkpoint integrity was monitored with fluorescently labeled ⍺-tubulin (GFP-Tub1 or mCherry-Tub1). Cells were imaged on agarose pads (2% agarose in SC medium + 2% glucose + 100 µM IAA) as described in Microscopy at 5 min interval for 4 to 5 hours. Images were analyzed by scoring the following 4 categories: spindle elongated in the mother cell and the cell was arrested in anaphase with elongated spindle (1), spindle elongated in the mother cell and the cell exited mitosis with spindle disassembled in the mother cell (2), spindle elongated in the mother cell but later extended into the bud and exited mitosis with one spindle pole in the bud (3), and spindle elongated into the bud and exited mitosis with one spindle pole in the bud (4). Only categories 1&2 were used to calculate the percentages of cells arrested vs exited in mother.

### Modeling Tem1’s nucleotide cycle

Virtual Cell software (VCell^47,48^) version 7.5.0 was used to model Tem1’s nucleotide cycle. The model, “Tem1_Bfa1” by user “xiaoxuez”, can be accessed within the VCell software (available at https://vcell.org). Two compartments (cytosol and SPB) were set up for the GAP stimulated GTP hydrolysis and GEF-independent nucleotide exchange where the GAP (represented in the model as “Bfa1”) was placed at SPB to simulate anaphase cells. We have also simulated metaphase cells by distributing the GAP evenly between the cytosol and SPB and found no difference in the results. Each simulation started with 25 nM Tem1^GTP^ (∼ 1500 molecules/cell) in the cytosol and ran for 300 s to reach steady state. The parameters used can be found in the online model and the parameters scanned are summarized in Fig. S7B.

---

**Figure S1:**
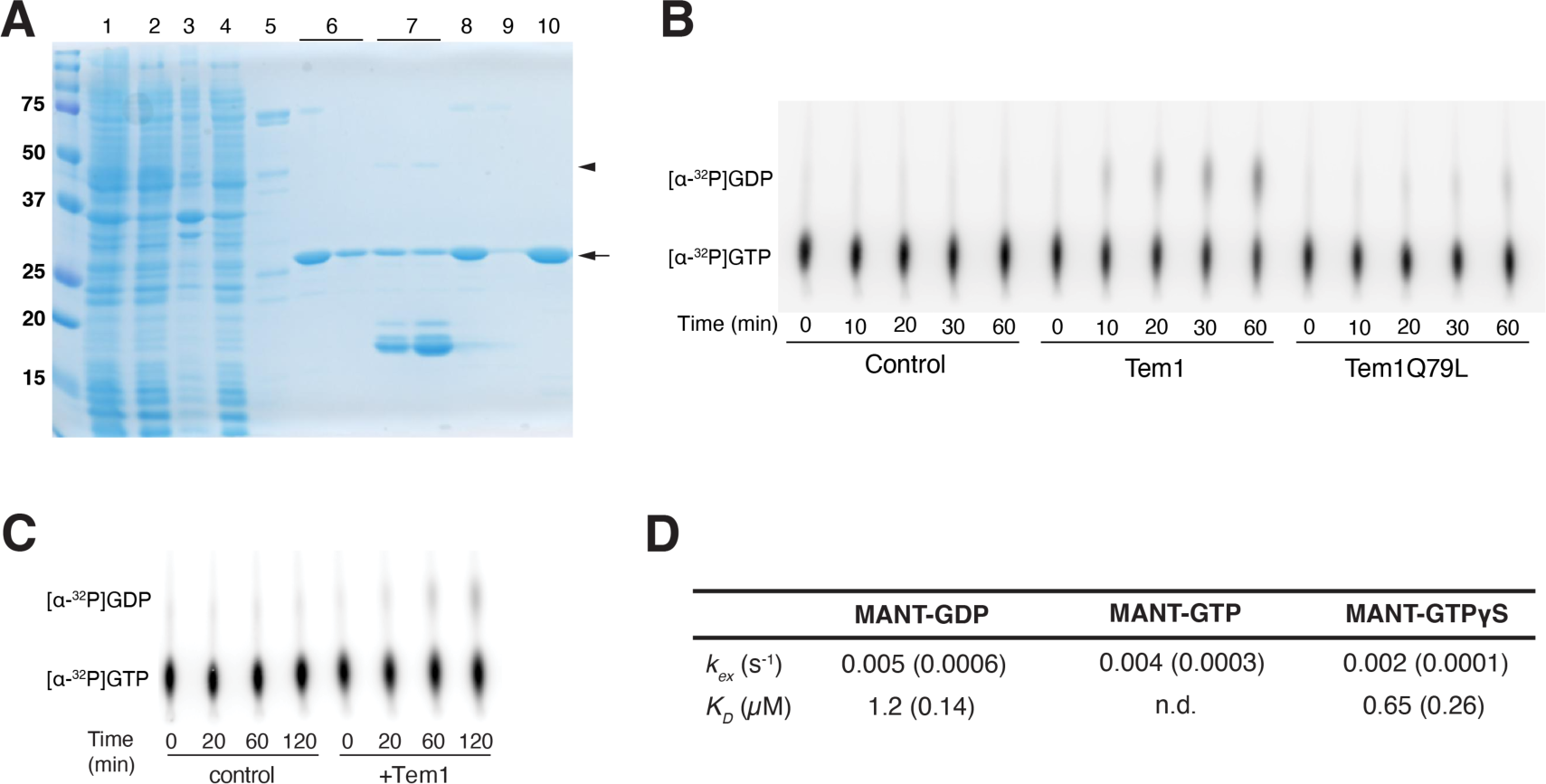
Biochemical characterization of Tem1. (**A**) Purification of Tem1. Tem1 was expressed in *E. coli* as a 14xHis-SUMO-Tem1 fusion with p2579. *Lane 1*, lysate; *lane 2*, soluble fraction; *lane 3*, insoluble fraction; *lane 4*, flowthrough of Ni-NTA column; *lane 5*, wash with lysis buffer; *lane 6*, elution after overnight digest with SENP protease which contains cleaved Tem1; *lane 7*, elution with high imidazole buffer where the remaining 14xHis-SUMO and uncut fusion protein were eluted; *lane 8*, input for Mono S column; *lane 9*, wash with buffer A; *lane 10*, combined peak elution of Tem1 from Mono S. Arrowhead marks 14xHis-SUMO-Tem1 fusion (41 kDa) and arrow marks Tem1 (27 kDa). (**B-C**) Representative raw thin layer chromatography (TLC) results for measuring GTPase activity of recombinantly purified Tem1 and the hydrolysis deficient mutant Tem1Q79L. No protein was added to the control sample (buffer only). (**D**) Summary of biochemical properties of Tem1 *in vitro*. Shown are the averages of three independent experiments and SD in parentheses. n.d., not determined (due to GTP hydrolysis). The difference of the observed exchange rate between MANT-GTP (*k_obs_* ≈ *k_off_* (GTP) + *k_hydrolysis_*) and MANT-GTPγS (*k_obs_* ≈ *k_off_* (GTPγS)) is likely not driven by GTP hydrolysis as previously assumed in filter-binding assays because GTP hydrolysis occurs at a rate an order of magnitude slower than dissociation (*k_obs_* ∼ 10^−3^ s^−1^, while *k_hydrolysis_* ∼ 10^−4^ s^−1^). Rather, we propose that the observed rates reflect differences in affinity or off rate of the nucleotides (*k_obs_* ≈ *k_off_*, when *k_hydrolysis_* << *k_obs_*). The higher affinity for MANT-GTPγS relative to MANT-GDP is consistent with the lower exchange rate for MANT-GTPγS we observed.

**Figure S2:**
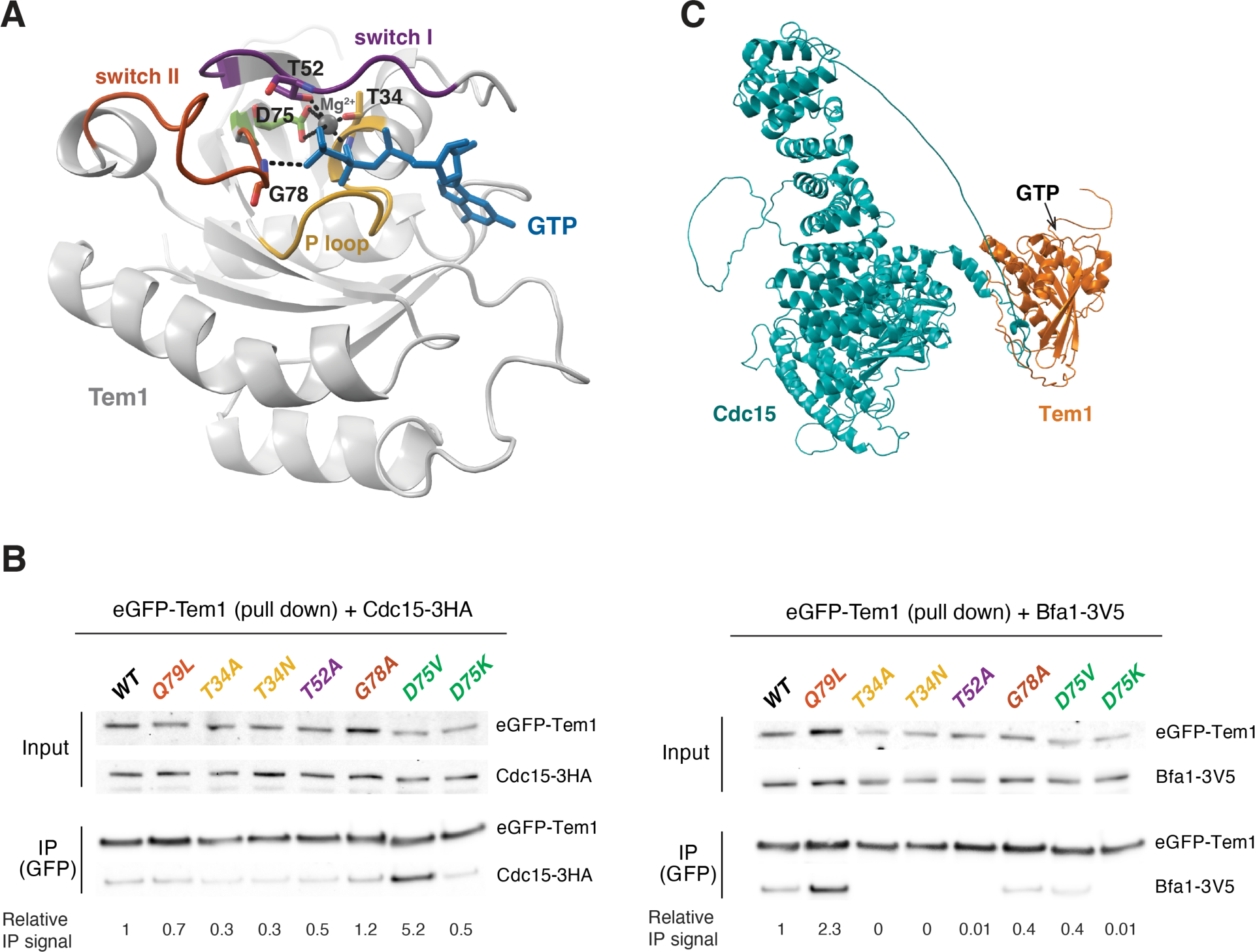
Interaction between Tem1 and Cdc15 or the GAP complex. (**A**) Mutated residues in Tem1. A model of Tem1 (residues 13–187 containing the core GTPase domain) was downloaded from Swiss Model (P38987_20210204) using Rab35 (GTP)-RUSC2 as a template. Tem1 backbone (gray ribbon), GTP (cyan), Mg^2+^ (dark gray sphere), P-loop with T34 (yellow), switch-I with T52 (purple), switch-II with G78 (orange) and D75 (green). T34, T52, and D75 coordinate the Mg^2+^ atom while G78 forms hydrogen bond with the γ-phosphate (dashed lines). T34 (Ras-S17) and D75 (Ras-D57) contribute to nucleotide binding and mutations in these residues have been shown to bias affinity toward GDP over GTP for Ras. T52 (Ras-T35) and G78 (Ras-G60) are invariant residues in switch I/II regions of small GTPases that form key contacts with the γ-phosphate in GTP to stabilize the active confirmation that is used by regulatory proteins and effectors to ‘sense’ the nucleotide status of the small GTPases. (**B**) Interaction between different Tem1 mutants and Cdc15 or the GAP complex via co-immunoprecipitation. yEGFP-Tem1 was immunoprecipitated from exponentially grown cells of y2885/y2886/y3209/y3210/y3208/y3211/y3212/y3213 (Cdc15-3HA with indicated yEGFP tagged Tem1 mutants) or y3452/y3472/y3457/y3458/y3456/y3455/y3459/y3460 (Bfa1-3V5 with indicated yEGFP tagged Tem1 mutants) and the presence of Cdc15-3HA or Bfa1-3V5 were analyzed by Western blot analysis. (**C**) AlphaFold2 predicted structure of Tem1-Cdc15 complex (DOI: 10.5452/ma-bak-cepc-0244).

**Figure S3:**
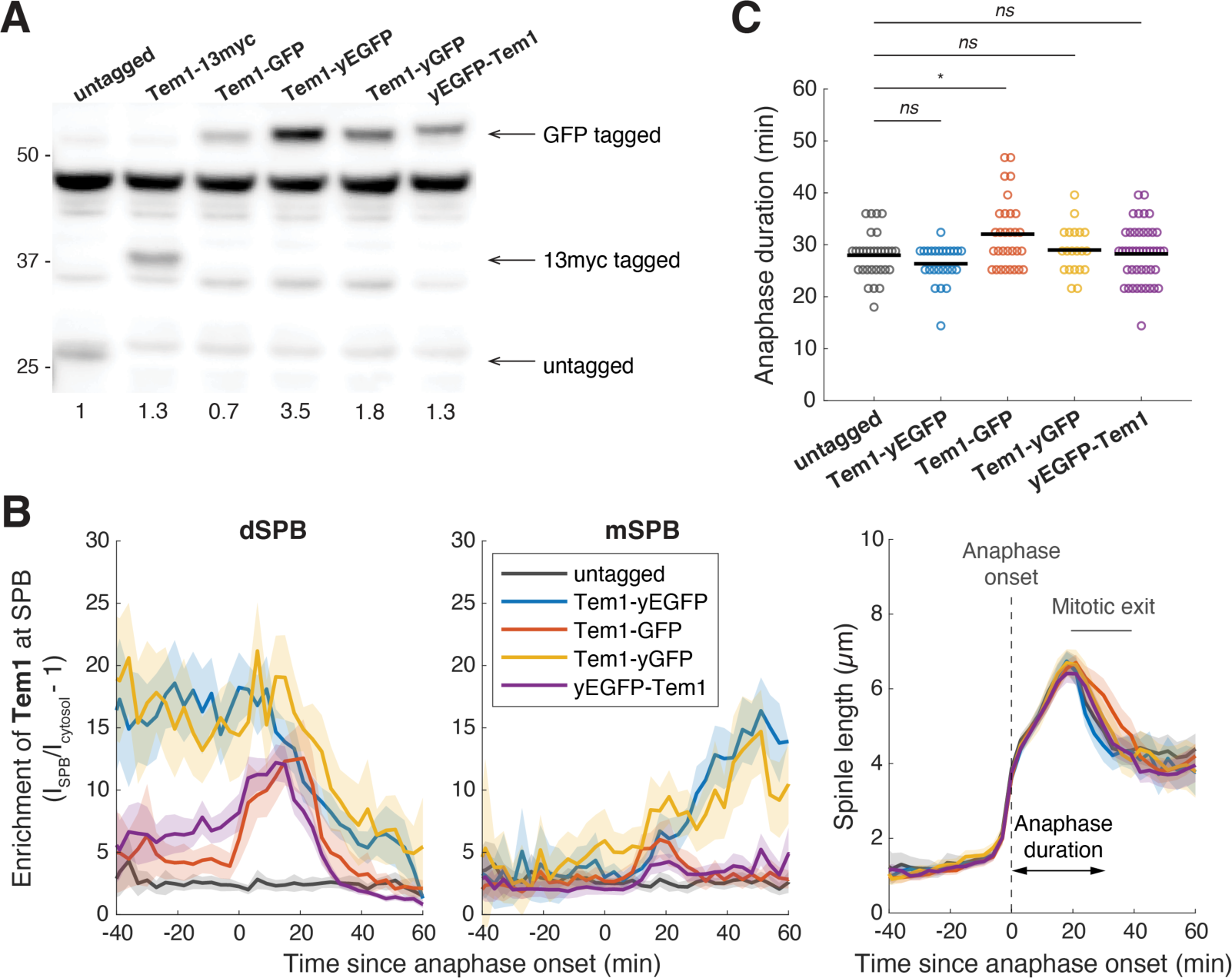
N-terminal fusion has a minimal perturbation on Tem1. (**A**) Expression level of different Tem1 fusions. Whole-cell lysates of exponentially growing yA2587 (untagged), yA1828 (Tem1-13myc), yA22523 (Tem1-GFP), yA21089 (Tem1-yEGFP), y1724 (Tem1-yGFP), and y1735 (yEGFP-Tem1) were analyzed by western blot with serum against Tem1. Cells were grown in YPD at 30 °C. C-terminal fusions of Tem1 stabilize the protein (unless a non-codon optimized GFP fusion is used). N-terminal fusion yEGFP-Tem1 is expressed at a level similar to wild-type Tem1 (1.3x) and cells expressing this fusion exit from mitosis at a rate similar to untagged cells (C). (**B**) SPB localization and spindle kinetics for different Tem1 fusions (y1008, y1267, y1271, y1739, and y1748; *n* = 39, 35, 39, 28, and 62 cells respectively) measured using the SPB marker Spc42-mScarlet-I. Cells were grown at 25 °C in SC medium + 2% glucose and imaged every 3 minutes for 4 hours. Solid lines represent the mean, shaded areas represent 95% confidence intervals. *TEM1-yEGFP* (codon-optimized eGFP) is known to be hyperactive (SPoC defective) whereas *TEM1-GFP* (not codon-optimized for yeast) is hypoactive as cells expressing Tem1-GFP delays exit from mitosis for more than 3 minutes. Tem1-yGFP (codon-optimized GFP) behaves more like Tem1-yEGFP rather than Tem1-GFP for SPB localization indicating it is the codon usage, likely due to changes in expression level as shown in (A), that is responsible for the difference in activity and localization profiles. (**C**) Anaphase duration of the cells shown in (B). Solid lines represent the median. *ns*, not significant (*p* > 0.05), \**p* < 0.05 by two-sided Wilcoxon rank sum test.

**Figure S4:**
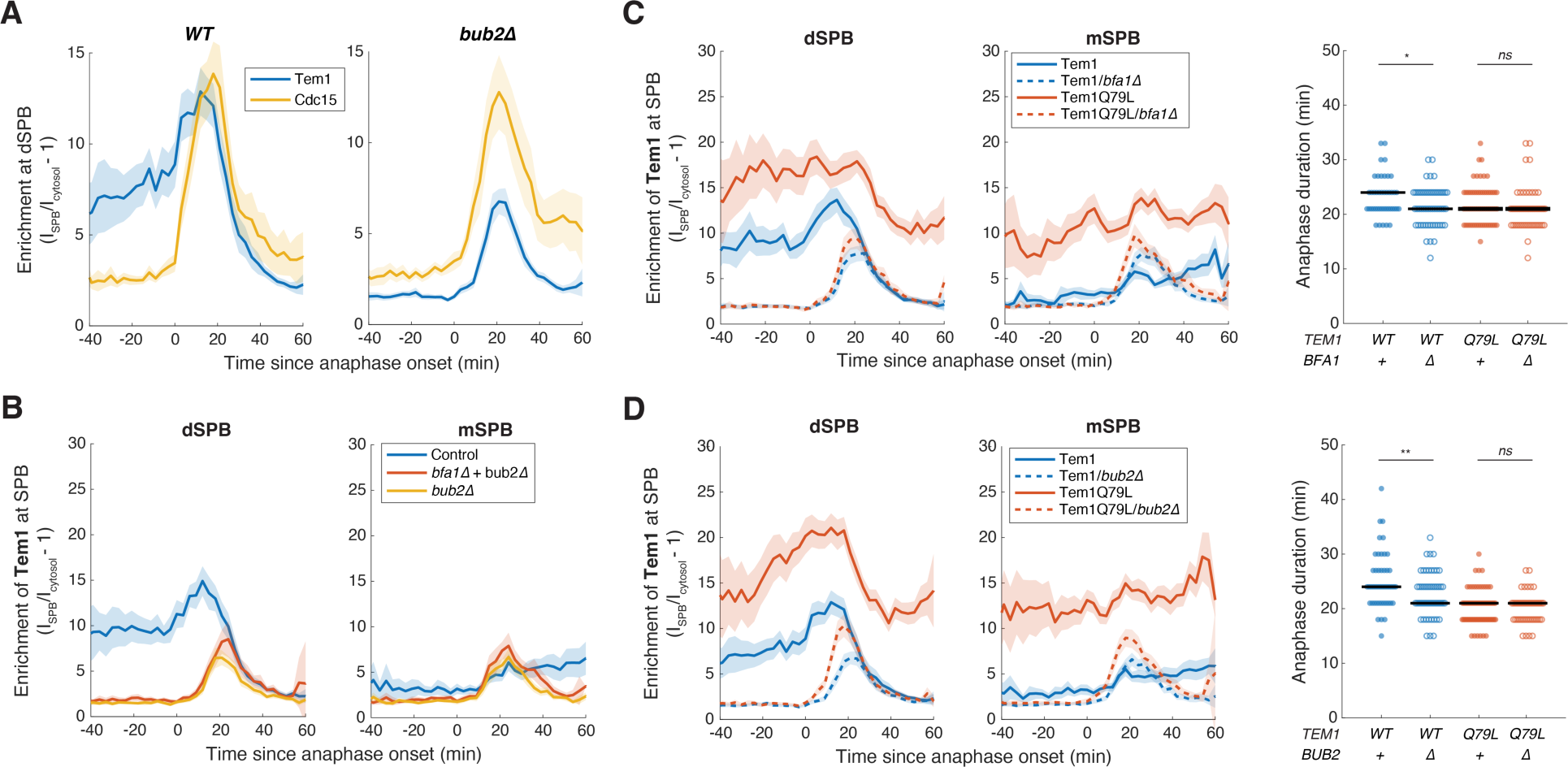
Localization of Tem1 to the SPB depends on the GAP complex Bub2-Bfa1. (**A**) Comparison of SPB localization for Tem1 and Cdc15 in the presence or absence of *BUB2*. (**B**) SPB localization of Tem1 with or without *BFA1* & *BUB2*. Cells of y2015 (Tem1, *n* = 44 cells), y3717 (Tem1 in *bfa1Δ+bub2Δ*, *n* = 42 cells), and y3718 (Tem1 in *bub2Δ*, *n* = 50 cells) were grown at 25 °C in SC medium + 2% glucose and imaged every 3 minutes for 4 hours. Solid lines represent the average. Shaded areas represent 95% confidence intervals. (**C-D**) SPB localization of Tem1 or the GTP-locked Tem1Q79L with or without the GAP component *BFA1* (C) or *BUB2* (D) and the anaphase duration for the corresponding cells. Cells of y2015 (Tem1, *n* = 54 cells), y2098 (Tem1 in *bfa1Δ*, *n* = 64 cells), y2019 (Tem1Q79L, *n* = 79 cells), y2133 (Tem1Q79L in *bfa1Δ*, *n* = 64 cells), and y1748 (Tem1, *n* = 65 cells), y1929 (Tem1 in *bub2Δ*, *n* = 72 cells), y1824 (Tem1Q79L, *n* = 91 cells), y1930 (Tem1Q79L in *bub2Δ*, *n* = 64 cells) were grown at 25 °C in SC medium + 2% glucose and imaged every 3 minutes for 4 hours. Solid lines represent the median. *ns*, not significant (*p* > 0.05), \**p* < 0.05 by two-sided Wilcoxon rank sum test.

**Figure S5:**
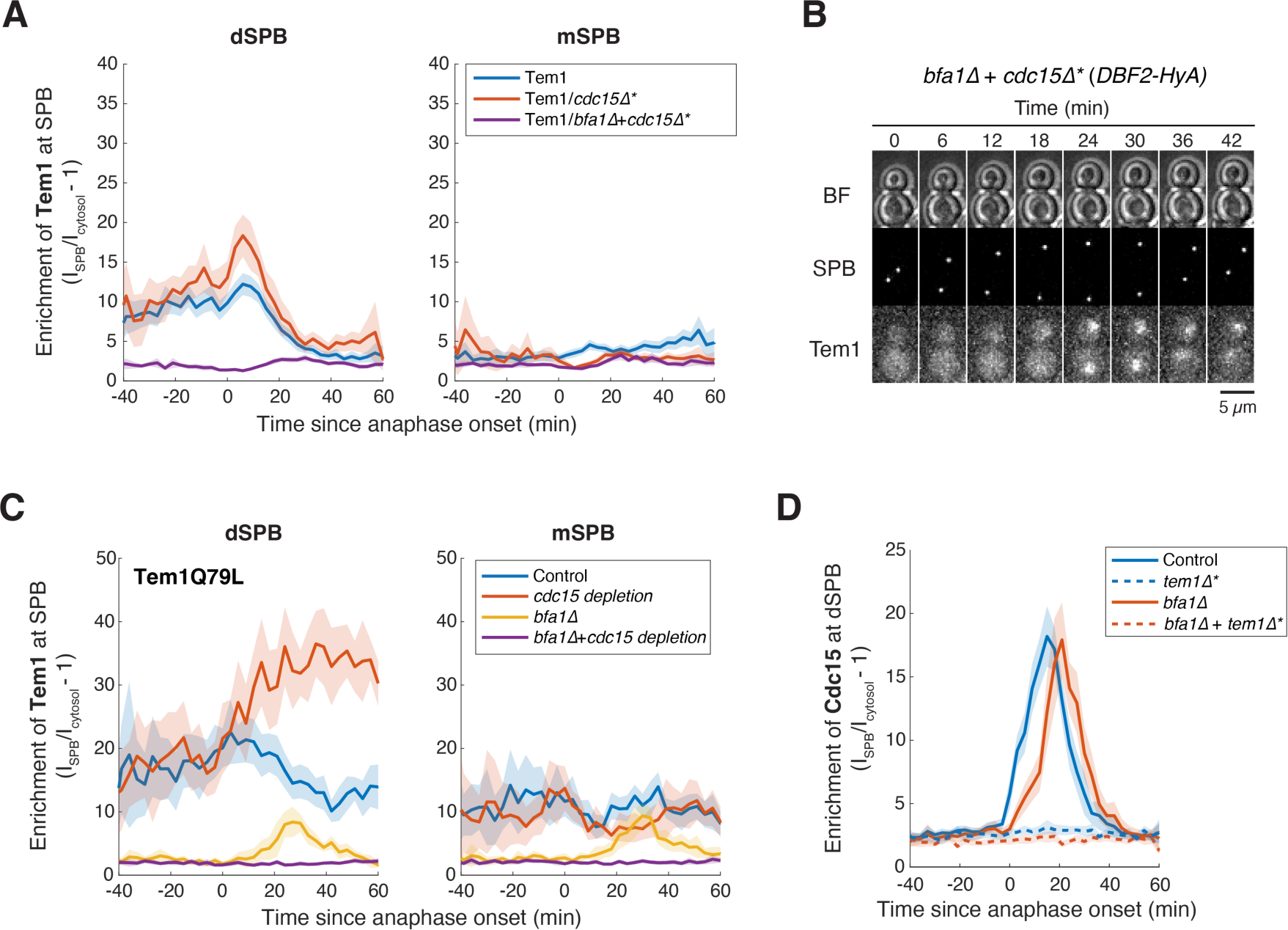
Localization of Tem1 to the SPB depends on Cdc15. (**A**) SPB localization of Tem1 with or without *BFA1*&*CDC15*. Cells of y2898 (Tem1, *n* = 61 cells), y2910 (Tem1 in *cdc15Δ*, *n* = 24 cells), and y2899 (Tem1 in *bfa1Δ+cdc15Δ*, *n* = 42 cells) were grown at 25 °C in SC medium + 2% glucose and imaged every 3 minutes for 4 hours. * denotes the presence of the hyperactive *DBF2-HyA* which rescues *cdc15Δ*. (**B**) Representative localization of Tem1 in *bfa1Δ+cdc15Δ* cell kept alive with *DBF2-HyA* (y2899). No SPB localization was observed but nuclear import and degradation (Fig. S12) still occurred contributing to the slight increase in SPB enrichment observed in late anaphase (A). (**C**) SPB localization of Tem1Q79L with or without depleting Cdc15 (*CDC15-AID* +/− *osTIR*) in the presence or absence of *BFA1*. Cells of y3231 (Tem1Q79L control, *n* = 18 cells), y3226 (Tem1Q79L with Cdc15 depletion, *n* = 14 cells), y3230 (Tem1Q79L in *bfa1Δ*, *n* = 16 cells), and y3225 (Tem1Q79L in *bfa1Δ* with Cdc15 depletion, *n* = 17 cells) were grown at 25 °C in SC medium + 2% glucose + 200 μM auxin IAA and imaged every 3 minutes for 4 hours. (**D**) SPB localization of Cdc15 still depends on Tem1 in the absence of the GAP. Cells of y1275 (Cdc15-yEGFP, *n* = 75 cells), y2945 (Cdc15-yEGFP in *tem1Δ*, *n* = 41 cells), y2958 (Cdc15-yEGFP in *bfa1Δ*, *n* = 37 cells), and y3010 (Cdc15-yEGFP in *tem1Δ*+*bfa1Δ*, *n* = 37 cells) were grown at 25 °C in SC medium + 2% glucose and imaged every 3 minutes for 4 hours. Cells with *tem1Δ* were kept alive with *DBF2-HyA*.

**Figure S6:**
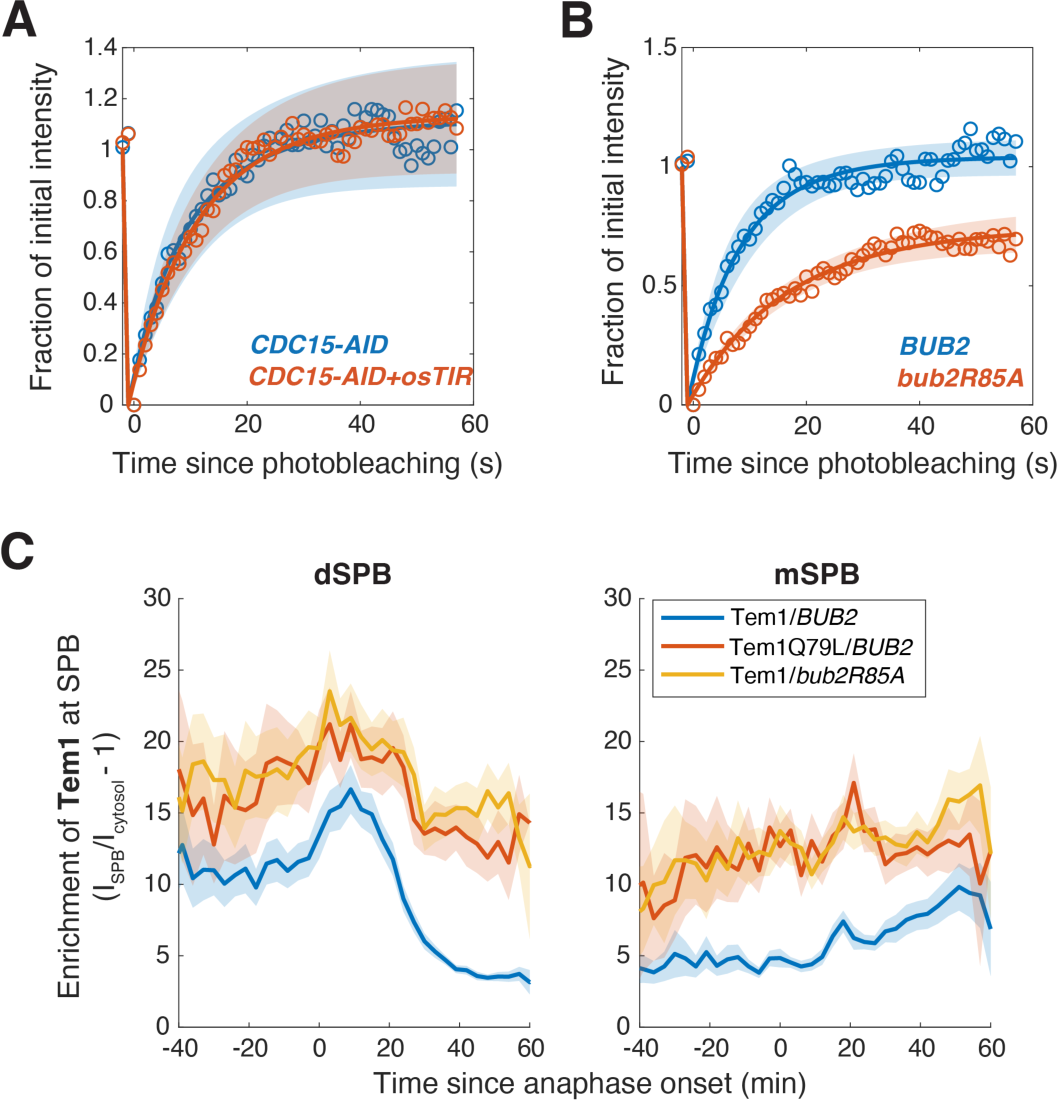
GAP stimulated GTP hydrolysis drives the fast turnover of Tem1 at the dSPB. (**A**) FRAP analysis of Tem1 at dSPB in anaphase with or without depleting Cdc15. Cells of y3901 (*CDC15-AID*, *n* = 6 cells) and y3900 (*CDC15-AID+OsTIR*, *n* = 9 cells) were grown and imaged at room temperature in SC medium + 2% glucose + 100 μM IAA. Circles represent the average measurements of double normalized fluorescence intensities after correcting for photo-bleaching during acquisition. Solid lines are the average fit and shaded areas represent SD. (**B**) FRAP analysis of Tem1 at dSPB in anaphase with hydrolysis defective Bub2 mutant (*bub2R85A*). Cells of y2015 (*BUB2*, *n* = 6 cells) and y2971 (*bub2R85A*, *n* = 6 cells) were grown and imaged at room temperature in SC medium + 2% glucose. Circles represent the average measurements of double normalized fluorescence intensities after correcting for photo-bleaching during acquisition. Solid lines are the average fit and shaded areas represent SD. (**C**) SPB localization of Tem1 in cells with defective GTP hydrolysis. Cells of y2015 (*WT*, *n* = 69 cells), y2019 (*TEM1Q79L*, *n* = 42 cells), and y2971 (*bub2R85A*, *n* = 51 cells) were grown at 25 °C in SC medium + 2% glucose and imaged every 3 minutes for 4 hours. Solid lines represent the average. Shaded areas represent 95% confidence intervals.

**Figure S7:**
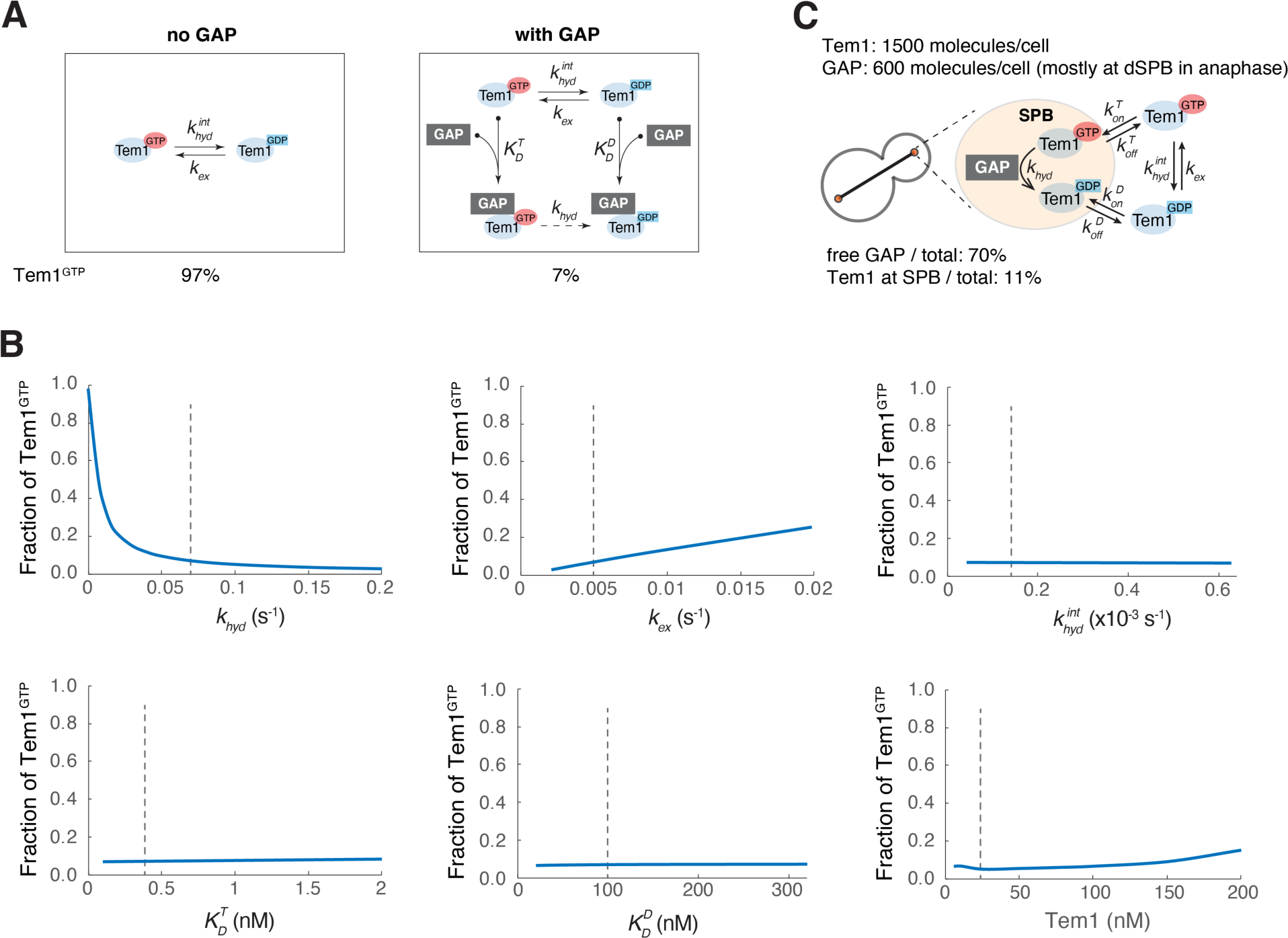
A simple model for Tem1’s nucleotide cycle. **(A)** A simplified model for Tem1’s GTP/GDP cycle with or without the GAP. **(B)** Fraction of Tem1^GTP^ as predicted by the model (with GAP) as a function of the GAP-stimulated GTP hydrolysis rate *k_hyd_*, Tem1’s intrinsic nucleotide exchange rate *k_ex_*, Tem1’s intrinsic GTP hydrolysis rate *k_hyd_^int^*, dissociation constant for GAP binding Tem1^GTP^ *K_D_^T^*, dissociation constant for GAP binding Tem1^GDP^ *K_D_^D^*, or Tem1 concentration in the cell. Dashed lines represent the values used to make the predictions shown in A. **(C)** A simplified model for Tem1’s GTP/GDP cycle at dSPB in anaphase. The same model parameters were used as in A.

**Figure S8:**
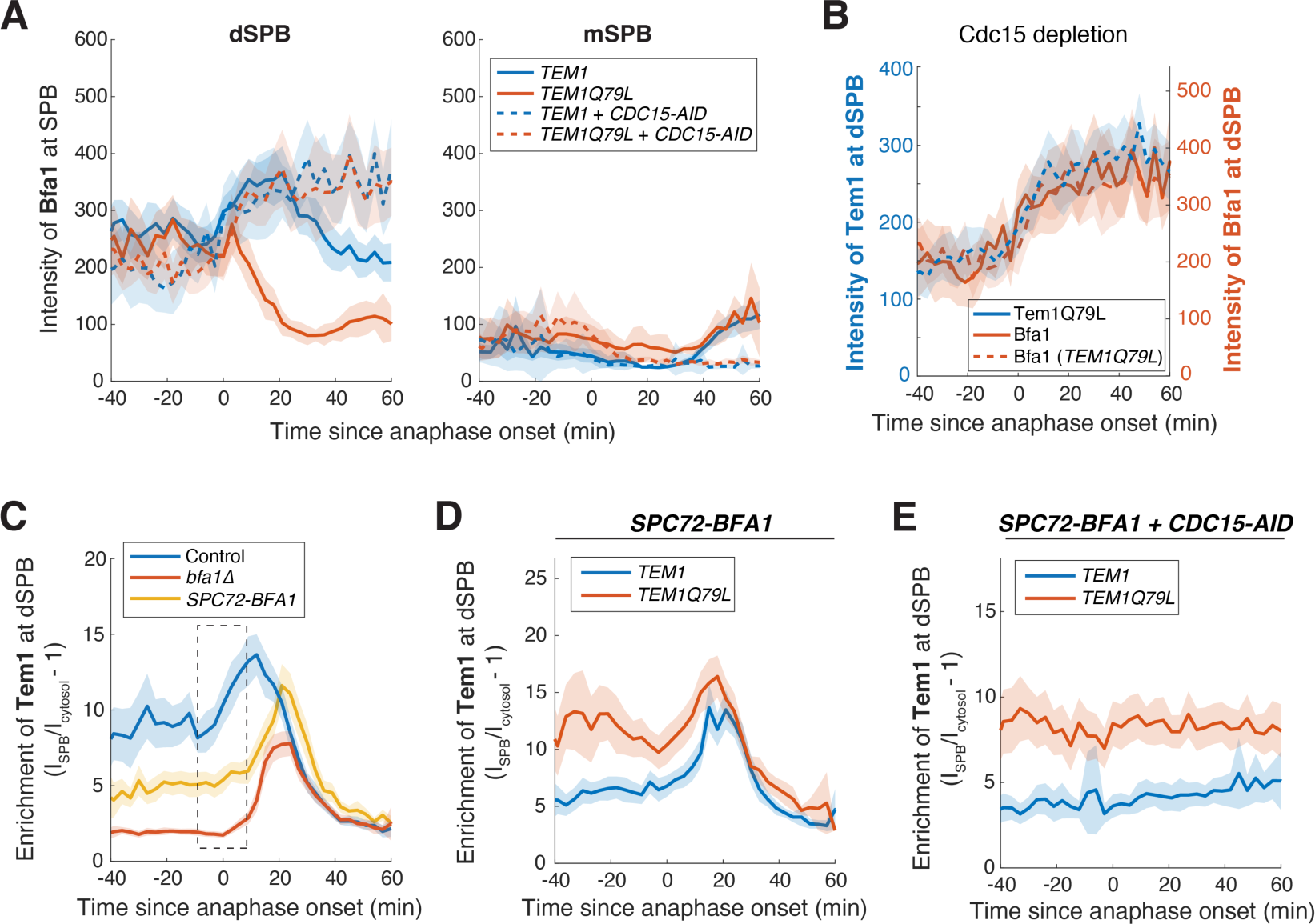
The GAP is redistributed upon anaphase onset. **(A)** dSPB localization of Bfa1 in cells expressing *TEM1* or *TEM1Q79L* with or without Cdc15 depleted. Cells of y1374 (*TEM1*, *n* = 42 cells), y2194 (*TEM1Q79L*, *n* = 30 cells), y4123 (*TEM1 + CDC15-AID*, *n* = 24 cells), and y4122 (*TEM1QL + CDC15-AID*, *n* = 32 cells) were grown at 25 °C in SC medium + 2% glucose + 100 μM IAA. The effect of *TEM1Q79L* on Bfa1 SPB localization depends on Cdc15. With Cdc15 depleted, Bfa1 localization to dSPB in cells expressing *TEM1Q79L* behaved the same as in cells with *TEM1*. **(B)** Comparison of dSPB localization for Tem1Q79L (same data as in Fig. 5E), Bfa1 in *TEM1* cells (same data as in A), and Bfa1 in *TEM1Q79L* cells (same data as in A) with Cdc15 depleted. **(C)** dSPB localization of Tem1 with or without GAP redistribution. Cells of y2015 (*BFA1*, *n* = 54 cells), y2098 (*bfa1Δ*, *n* = 64 cells), and y2990 (*SPC72-BFA1* which tethers the GAP to the SPB and prevents its redistribution, *n* = 48 cells) were grown at 25 °C in SC medium + 2% glucose. The increase in dSPB localization upon anaphase onset observed for Tem1 is driven by GAP redistribution. Tethering Bfa1 to the SPB with *SPC72-BFA1* abolished the increase upon anaphase onset (see boxed area). **(D)** dSPB localization of Tem1 or Tem1Q79L in cells with SPB-tethered GAP (*SPC72-BFA1*). Cells of y2990 (*TEM1*, *n* = 48 cells) and y2991 (*TEM1Q79L*, *n* = 54 cells) were grown at 25 °C in SC medium + 2% glucose. **(E)** dSPB localization of Tem1 or Tem1Q79L in cells with SPB-tethered GAP (*SPC72-BFA1*) and Cdc15 depleted (*CDC15-AID*). Cells of y3227 (*TEM1*, *n* = 53 cells) and y4106 (*TEM1Q79L*, *n* = 53 cells) were grown at 25 °C in SC medium + 2% glucose + 100 μM IAA. dSPB localization of Tem1 and Tem1Q79L remained steady when Bfa1 was tethered to the SPB in cells with Cdc15 depleted. For all plots, cells were imaged every 3 minutes for 4 hours. Lines represent the average. Shaded areas represent 95% confidence intervals.

**Figure S9:**
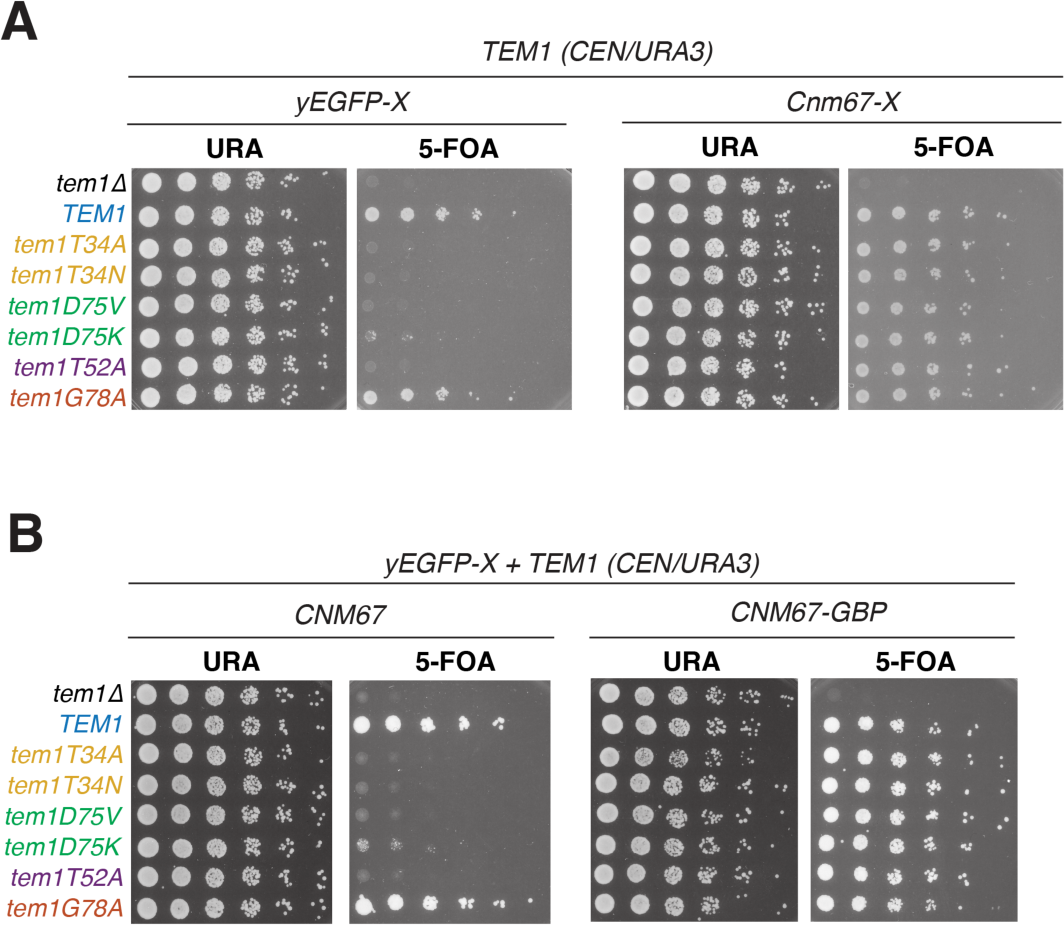
Tethering Tem1 to the SPB rescues the lethality of nucleotide binding mutants of Tem1. (**A**-**B**) Complementation analysis of Tem1 mutants with and without tethering to the SPB via direct fusion to Cnm67 (A) or using yGFP-Tem1 + Cnm67-GBP combo (B). 5-fold serial dilutions of strains y2891/y3202/y3203/y3204/y3206/y3207/y3214/y3205 (untethered) and y2891/y2893/y2927/y3187/y3190/y3191/y3189/y3188 (direct fusion as Cnm67-Tem1) or y3303/y3304/y3305/y3306/y3308/y3309/y3310/y3307 (with *CNM67-GBP*) were spotted onto plates with or without 5’-fluoroorotic acid (5-FOA) and incubated at 25°C for 3 to 4 days. The presence of 5-FOA selects cells that are viable after losing the *TEM1 (URA3/CEN)* plasmid. Mutations in T34, D75, and T52 render Tem1 inactive and severely reduce viability. In contrast, the switch-II effector mutant Tem1G78A is active.

**Figure S10:**
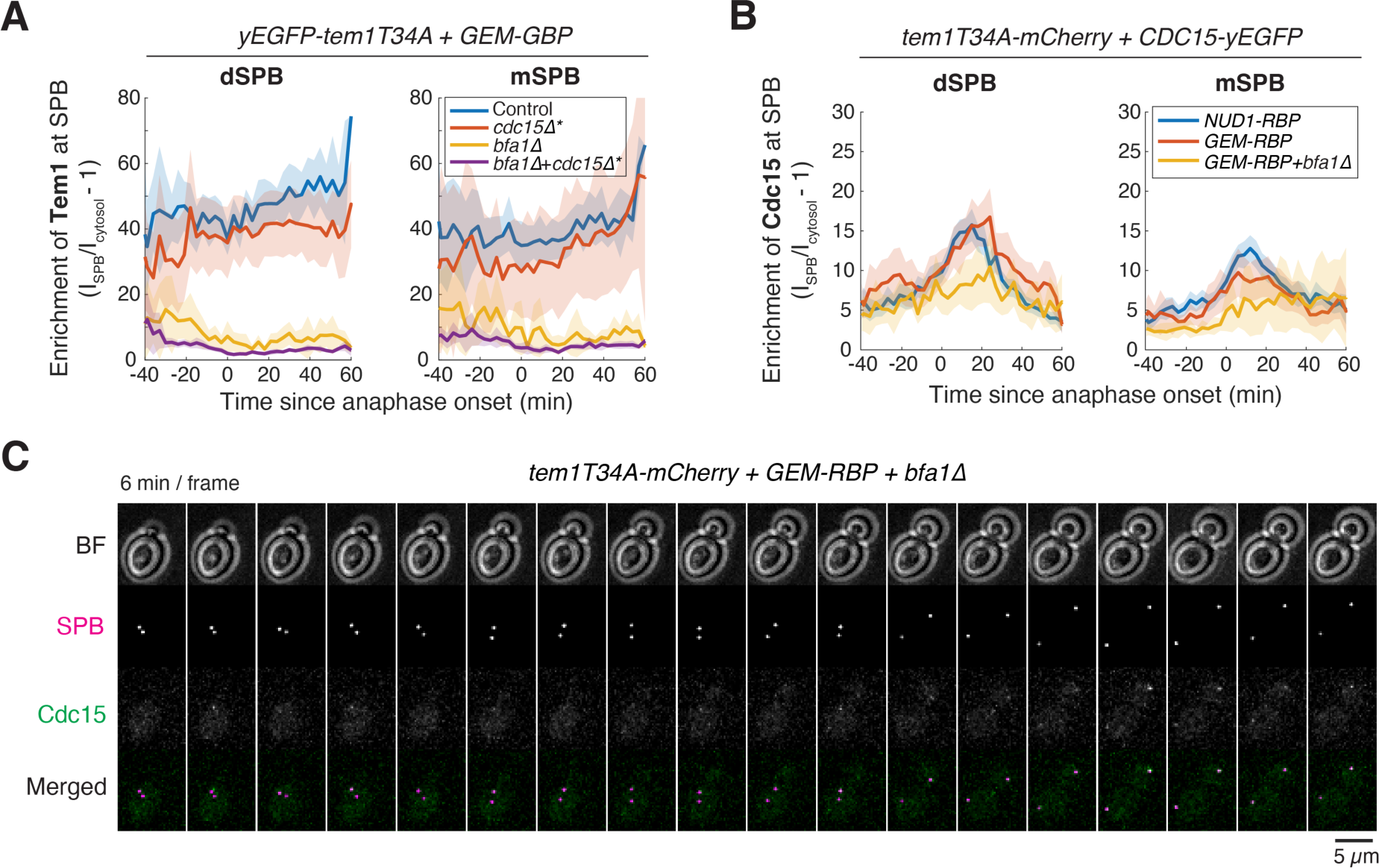
Concentrating Tem1 by tethering to GEMs activates the MEN. **(A)** SPB localization of GEM-tethered Tem1T34A in cells with or without *BFA1* or *CDC15*. Cells of y3630 (*BFA1+CDC15*, *n* = 27 cells), y3631 (*BFA1+cdc15Δ*, *n* = 18 cells), y3557 (*bfa1Δ+CDC15*, *n* = 21 cells), and y3510 (*bfa1Δ+cdc15Δ*, *n* = 21 cells) were grown at 25°C in SC medium + 2% glucose and imaged every 3 minutes for 4 hours. Solid lines represent the average. Shaded areas represent 95% confidence intervals. **(B)** SPB localization of Cdc15 in cells with SPB- or GEM-tethered Tem1T34A. Cells of y3585 (*NUD1-RBP*, *n* = 67 cells), y3589 (*GEM-RBP*, *n* = 35 cells), and y3588 (GEM-RBP+*bfa1Δ*, *n* = 14 cells) were grown at 25°C in SC medium + 2% glucose and imaged every 3 minutes for 4 hours. Solid lines represent the average. Shaded areas represent 95% confidence intervals. **(C)** Representative localization of Cdc15 in y3588.

**Figure S11:**
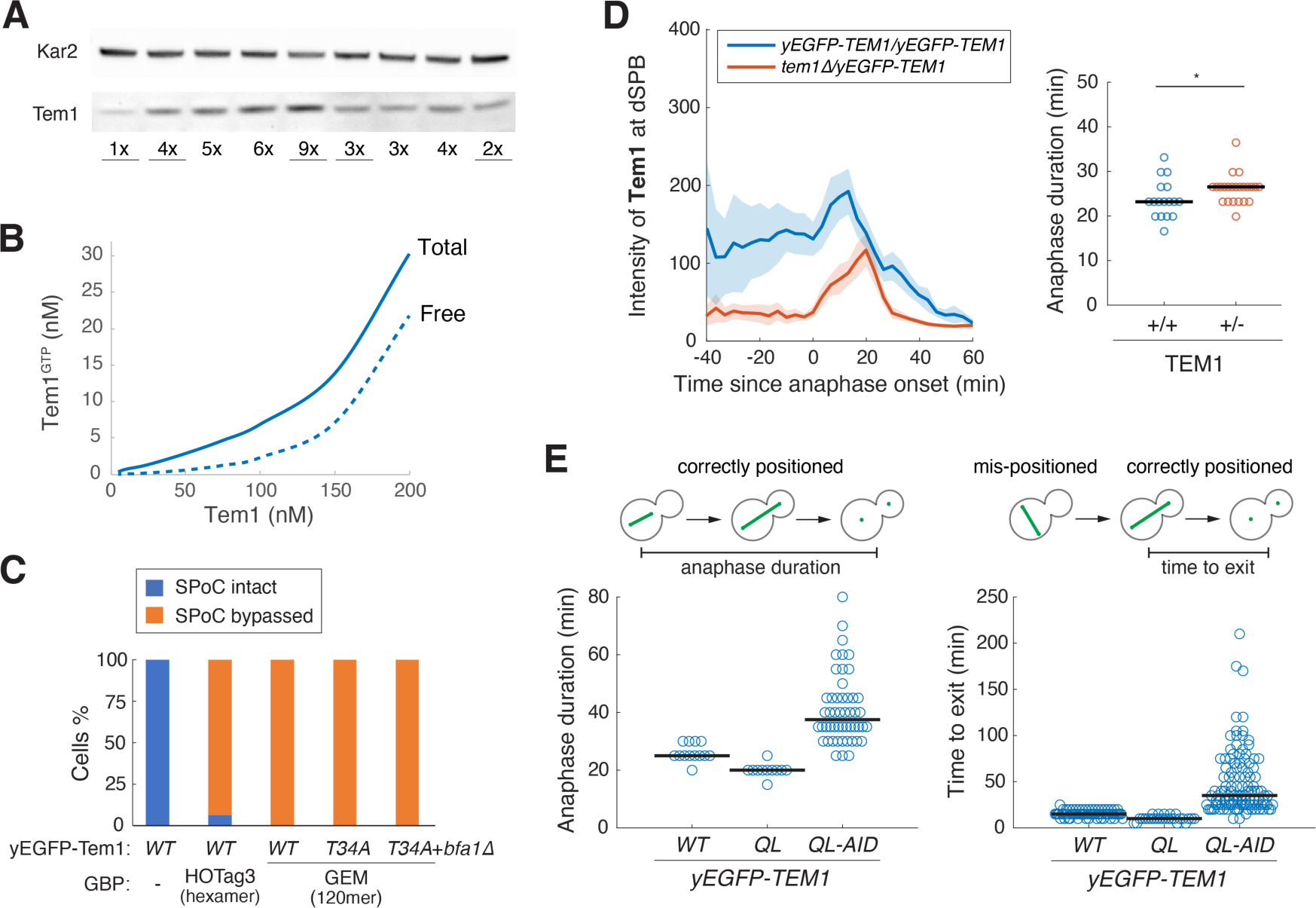
MEN activity is sensitive to Tem1 level. **(A)** Quantification of Tem1 expression level. Whole-cell lysates of exponentially growing cells with different copy numbers of Tem1 integrated at the *leu2* locus. Relative expression of Tem1 (indicated below) was quantified and normalized to the signal of Kar2. Underscored lanes represent strains used in this study: yA2587 (1xTem1, WT), y3693 (4xTem1), y3696 (9xTem1), y3692 (3xTem1), y3691 (2xTem1). **(B)** Predicted Tem1^GTP^ concentration as a function of Tem1 concentration in the cell (using the model illustrated in Fig. S7). **(C)** SPoC integrity in cells with locally clustered Tem1. Cells of y4097 (yEGFP-*TEM1*, *n* = 31 cells), y4096 (yEGFP-*TEM1 +* HOTag3-GBP, *n* = 63 cells), y4095 (yEGFP-*TEM1 +* GEM-GBP, *n* = 40 cells), y4100 (yEGFP-*tem1T34A +* GEM-GBP, *n* = 40 cells), and y4306 (yEGFP-*tem1T34A +* GEM-GBP + *bfa1Δ*, *n* = 45 cells) were grown at 25 °C in SC medium + 2% glucose + 100 μM IAA to induce spindle mispositioning by depleting Dyn1 and Kar9 and imaged every 5 minutes for 5 hours. Status of the spindle and cell cycle stages were monitored with mCherry-Tub1. **(D)** SPB localization of Tem1 (left) and distribution of anaphase duration (right) for diploid cells with two or one copy of *yEGFP-TEM1*. Cells of y3713 (two copies, *n* = 20 cells) and y3711 (one copy, *n* = 26 cells) were grown at 25°C in SC medium + 2% glucose and imaged every 3 minutes for 4 hours. Solid lines represent the average. Shaded areas represent 95% confidence intervals. \**p* < 0.05 by two-sided Wilcoxon rank sum test. Note that the SPB localization profile of Tem1 heterozygous deletion (∼50% expression level) resembles the hypomorphic Tem1-GFP (Fig. S3, ∼ 70% WT level). **(E)** Efficiency to exit from mitosis for GTP-locked Tem1 mutants with or without depletion. Same experiment as in Fig. 7E. Time to exit from mitosis as illustrated was measured for cells of y3817 (yEGFP-*TEM1*, *n* = 13 cells for anaphase duration and 72 cells for time to exit after correcting spindle position), y3818 (yEGFP-*TEM1Q79L*, *n* = 11 cells for anaphase duration and 27 cells for time to exit after correcting spindle position), and y4311 (yEGFP-*TEM1Q79L-AID\**, *n* = 52 cells for anaphase duration and 137 cells for time to exit after correcting spindle position).

**Figure S12:**
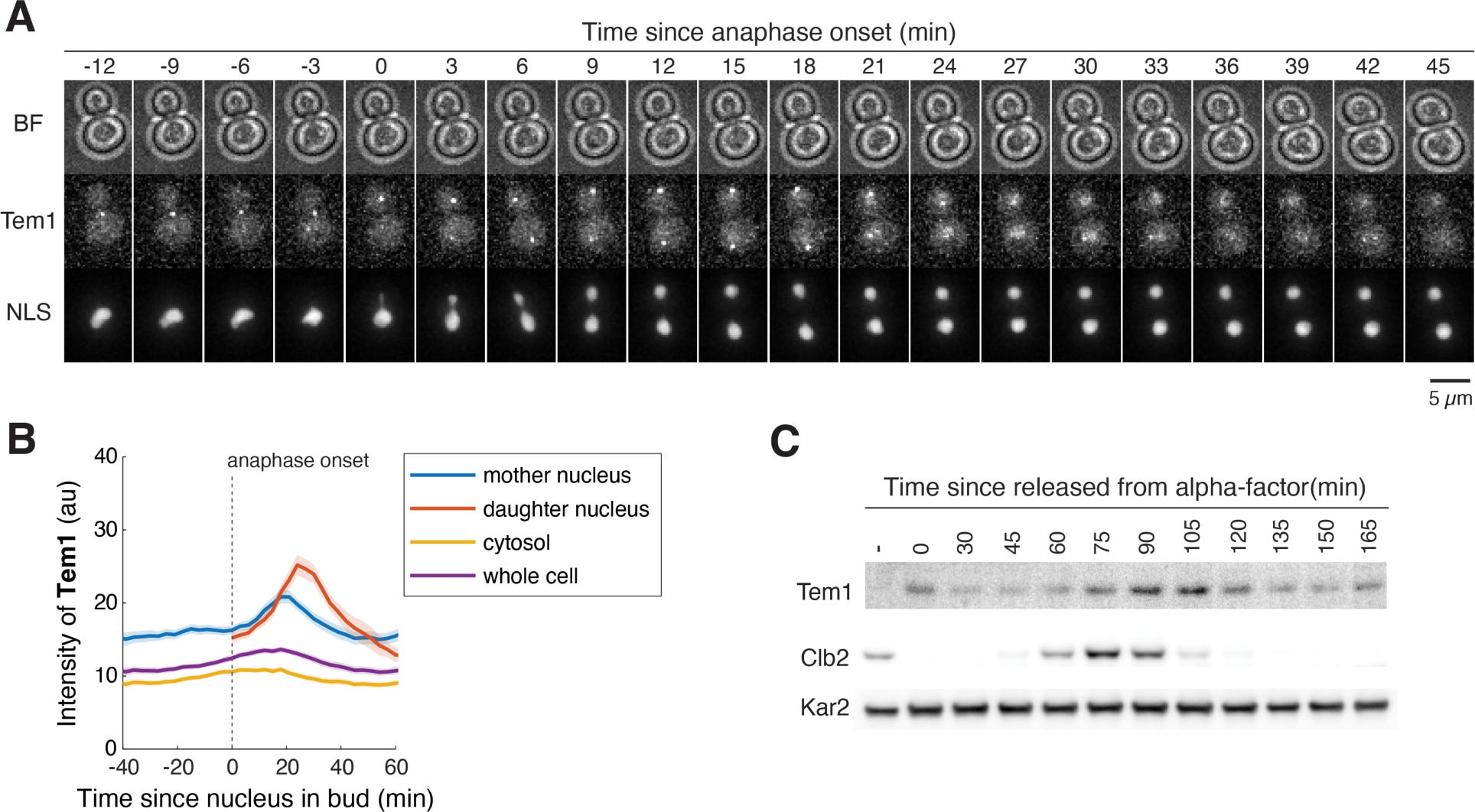
Tem1 level is cell cycle regulated. **(A)** Localization of Tem1 during the cell cycle where Tem1 is imported into the nucleus after exit from mitosis for degradation. y1879 (*yEGFP-TEM1* and *NLS_SV40_-2xmCherry*) cells were grown at 25 °C in SC medium + 2% glucose and imaged every 3 minutes for 4 hours. **(B)** Quantification of nuclear import and degradation of Tem1. Same cells as in (A). Lines represent the average; shaded areas represent 95% confidence intervals. **(C)** Tem1 level is cell cycle regulated. yA2587 (wild-type untagged strain) was arrested in G1 with α-factor (5 μg/ml) followed by release into YPD lacking pheromone. The amount of Tem1 protein were analyzed in whole-cell lysates by Western blot with anti-Tem1 serum. Clb2 was analyzed to determine when cells were in mitosis. Kar2 was used as internal loading control.

**Table S1.**
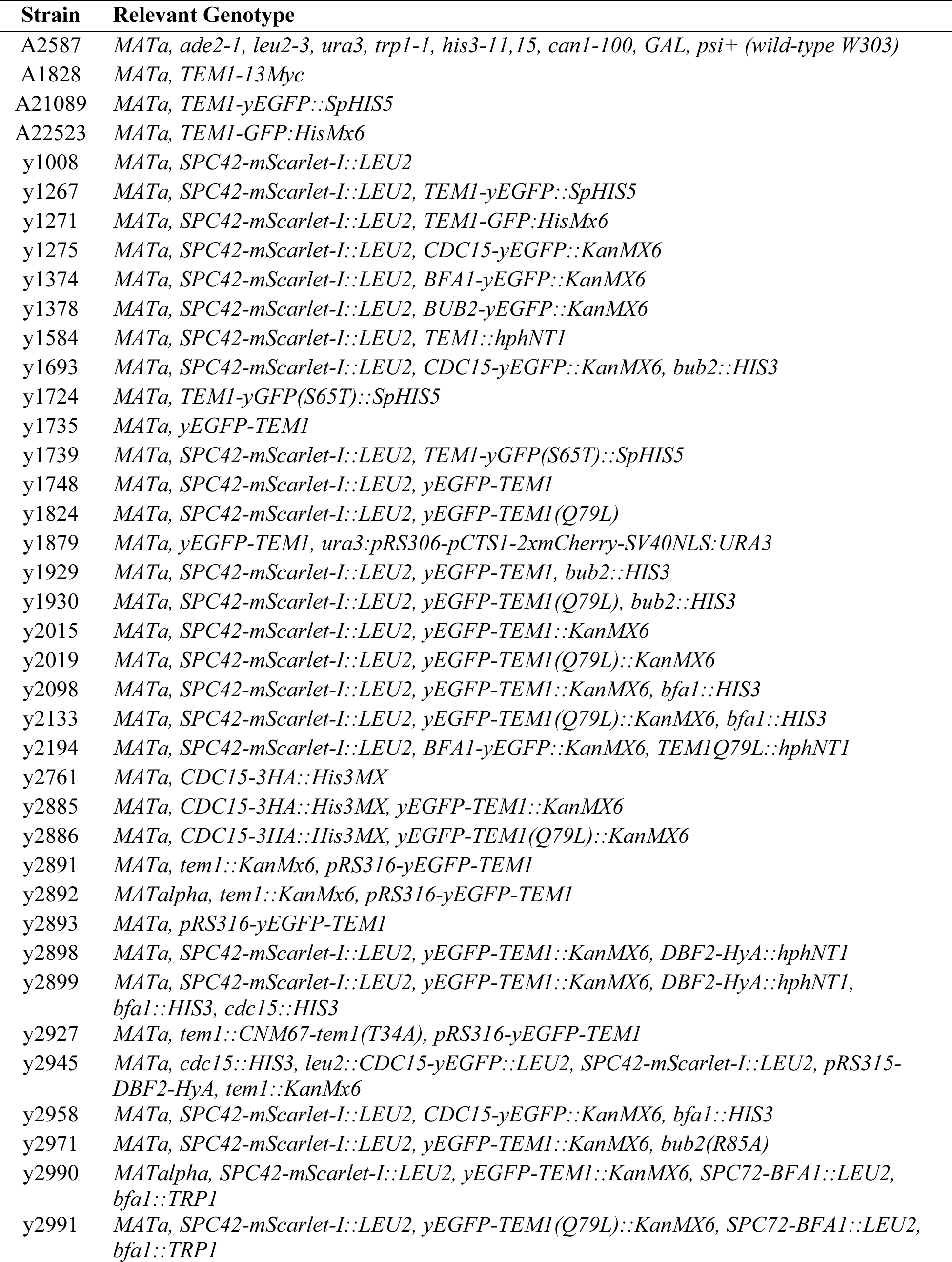
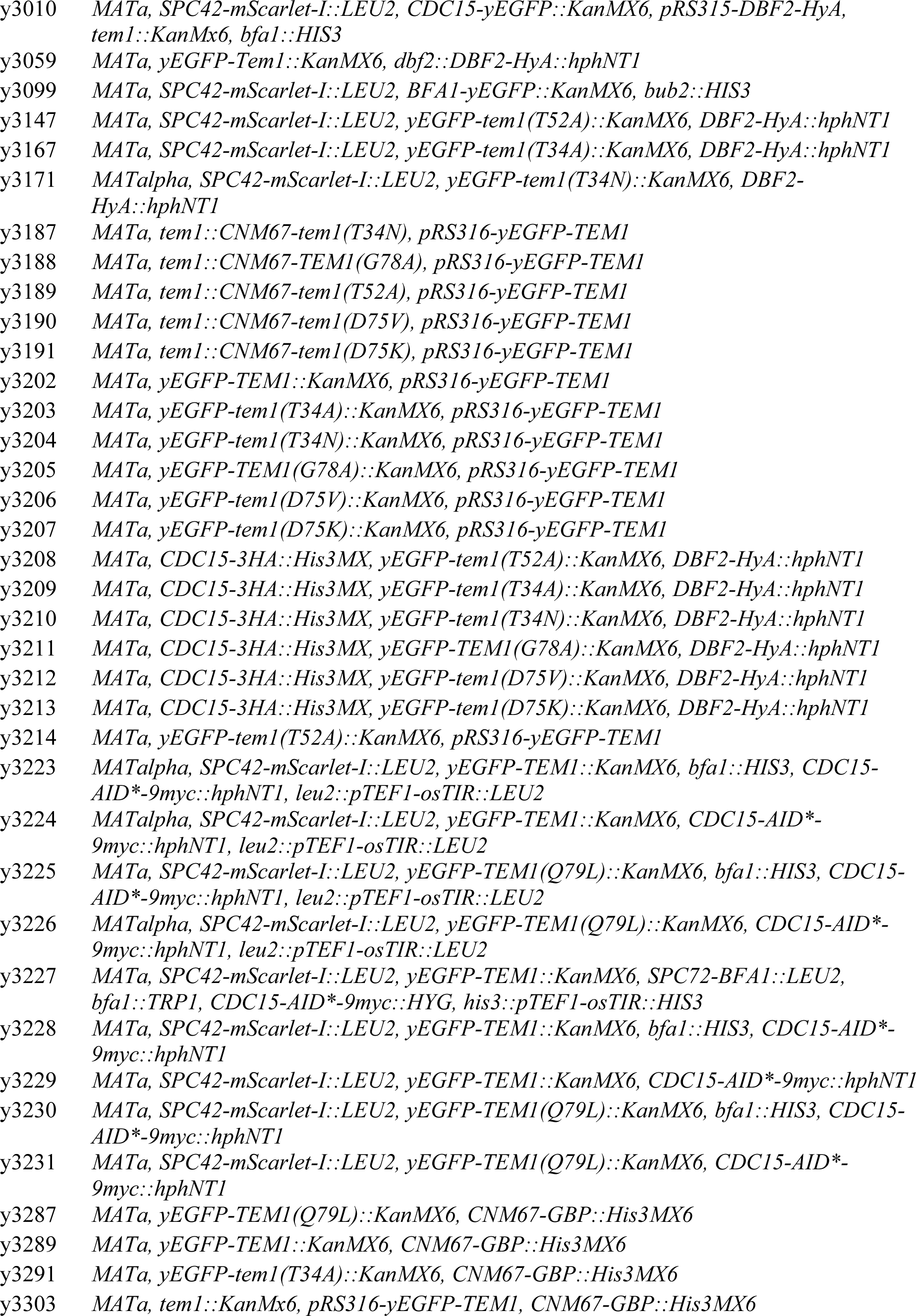
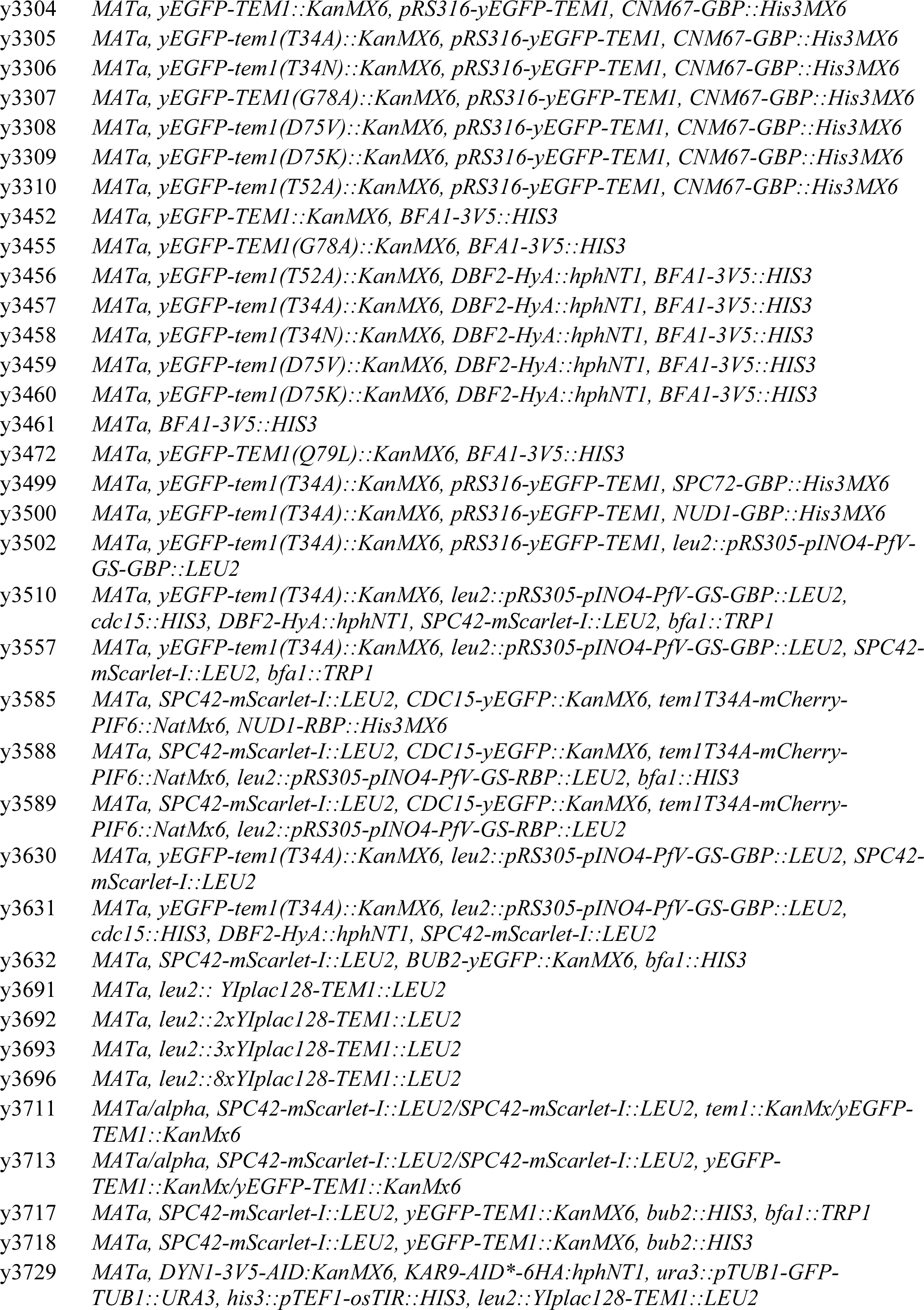
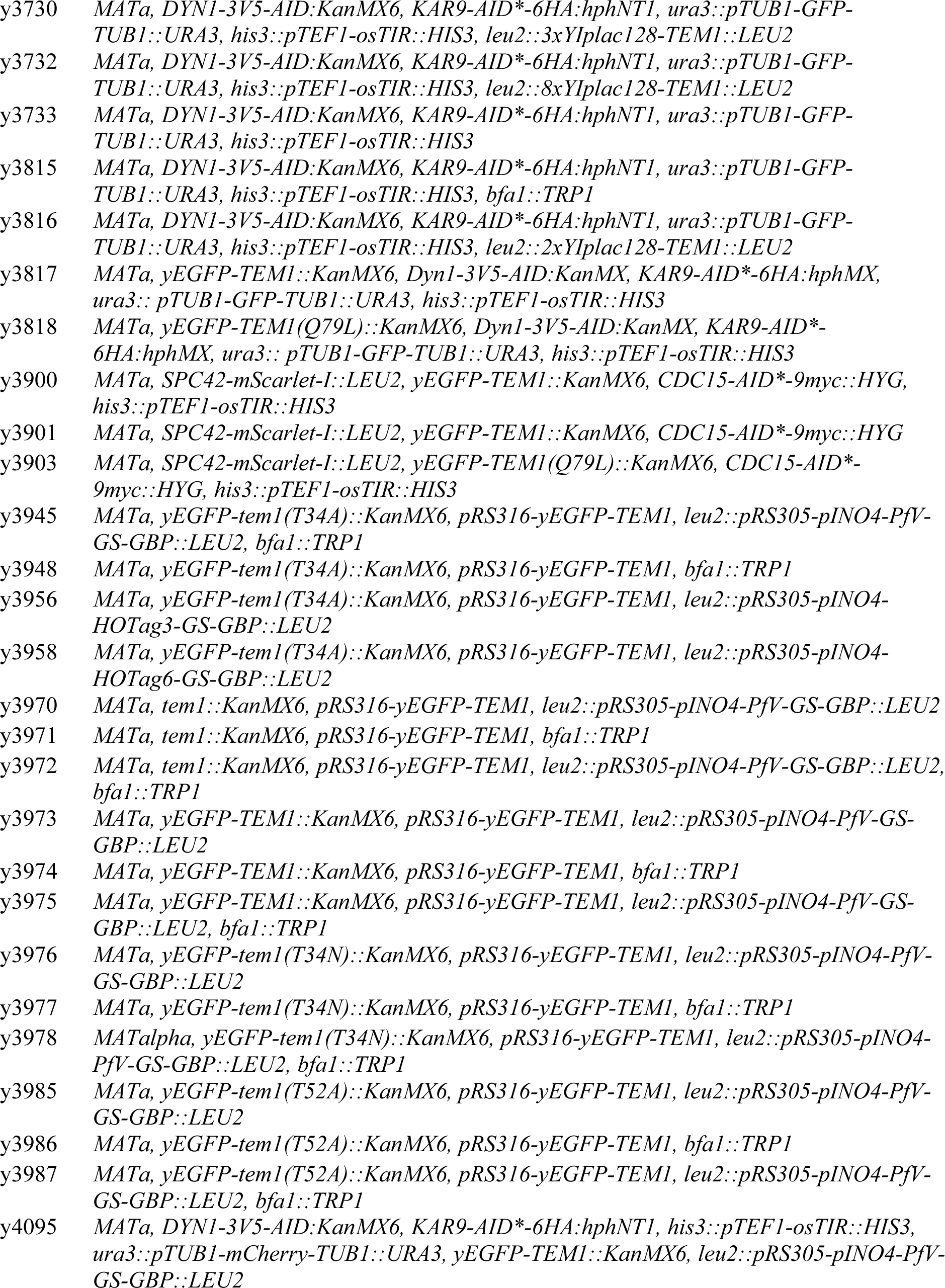
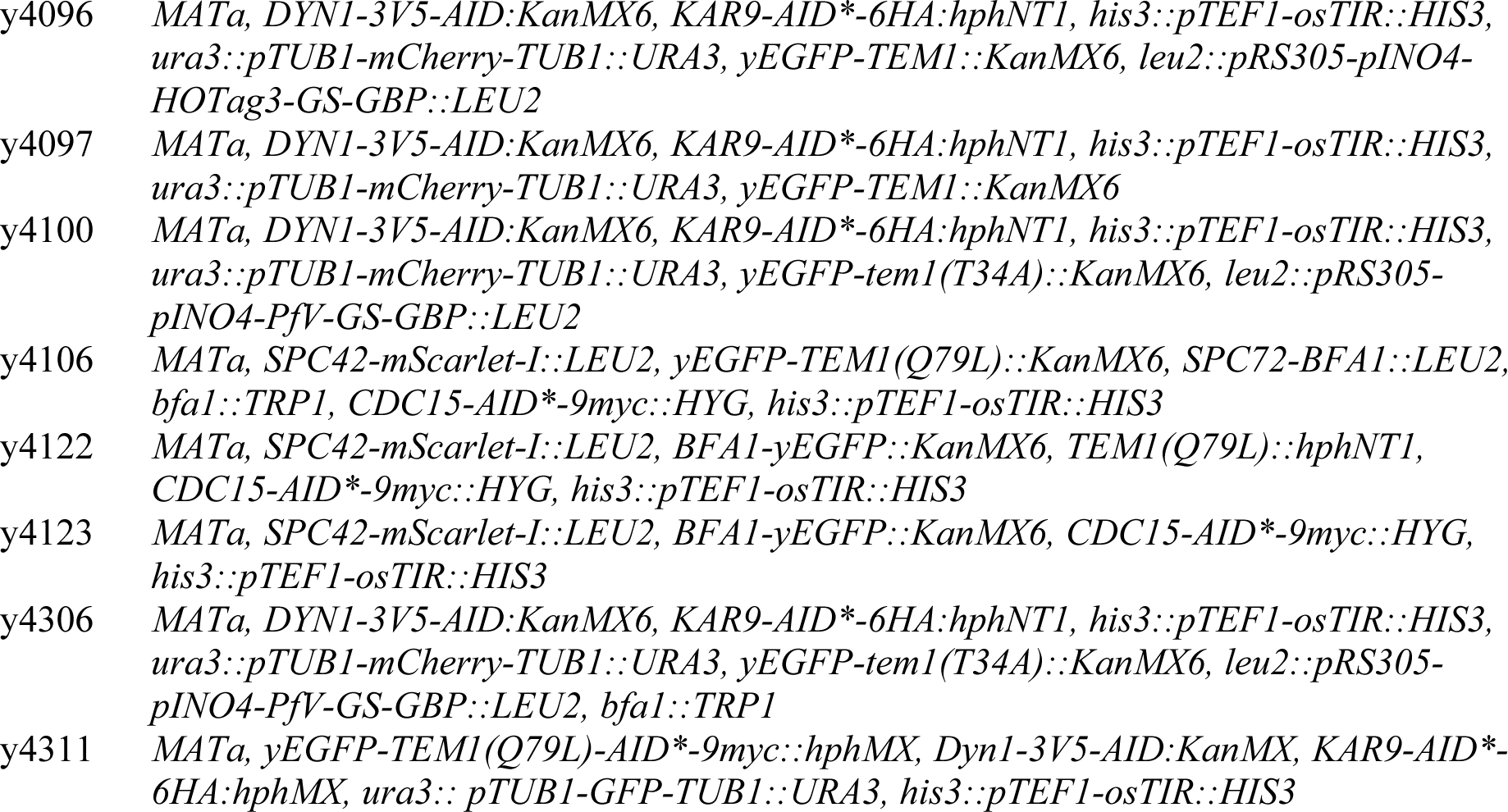
Yeast strains used in this study.

**Table S2.** Plasmids used in this study.

| **Plasmid** | **Description** | **Source** |
| --- | --- | --- |
| p998 | pFA6a-link-yEGFP-SpHIS5 | EUROSCARF |
| p1804 | pRS316-eGFP-Tem1 | This study |
| p2579 | pCOLADuet-1_14xHIS-SUMO-Tem1 | This study |
| p2717 | YIplac128-Tem1 (pSP596) | S. Piatti |
| p2718 | YIplac128-Tem1-Q79L (pSP597) | S. Piatti |
| p2816 | pFA6a-hphN-Dbf2-HyA | This study |
| p2868 | pFA6a-Vinnylinker-mScarlet-I-cgLEU2 (pUB1571) | E. Unal |
| p2888 | pFA6a-hphN-Tem1Q79L | This study |
| p2911 | pPGK1-Cas9-LEU (bRA90) | J. Haber |
| p2930 | pFA6a-linker-yGFP(S65T)-SpHIS5 | This study |
| p2938 | pRS305-pINO4-PfV-GS-Sapphire (pLH497) | L. Holt |
| p3009 | pFA6a-link-GBP-SpHIS5 | This study |
| p3010 | pFA6a-link-RBP-SpHIS5 | This study |
| p3011 | pRS305-pINO4-PfV-GS-GBP | This study |
| p3013 | pRS305-pINO4-PfV-GS-RBP | This study |
| p3021 | pCOLADuet-1-14xHIS-SUMO-Tem1Q79L | This study |
| p3034 | pDW363_AviTag-His6-Tem1_BirA | This study |
| p3045 | pRS305-pINO4-HOTag3-GS-GBP | This study |
| p3046 | pRS305-pINO4-HOTag6-GS-GBP | This study |

